# Hunger modulates exploration through suppression of dopamine signaling in the tail of striatum

**DOI:** 10.1101/2024.11.11.622990

**Authors:** Tarun Kamath, Bart Lodder, Eliana Bilsel, Isobel Green, Rochelin Dalangin, Paolo Capelli, Michelle Raghubardayal, Jessie Legister, Lauren Hulshof, Janet Berrios Wallace, Lin Tian, Naoshige Uchida, Mitsuko Watabe-Uchida, Bernardo L. Sabatini

**Affiliations:** Howard Hughes Medical Institute, Department of Neurobiology, Harvard Medical School, Boston, MA 02115, USA; Department of Molecular and Cellular Biology, Center for Brain Science, Harvard University, Cambridge, MA 02138, USA; CERVO Brain Research Center, Laval University, Quebec, QC G1E 1T2, Canada; Max Planck Florida Institute for Neuroscience, Jupiter, FL, 33458 USA

## Abstract

Caloric depletion leads to behavioral changes that help an animal find food and restore its homeostatic balance. Hunger increases exploration and risk-taking behavior, allowing an animal to forage for food despite risks; however, the neural circuitry underlying this change is unknown. Here, we characterize how hunger restructures an animal’s spontaneous behavior as well as its directed exploration of a novel object. We show that hunger-induced changes in exploration are accompanied by and result from modulation of dopamine signaling in the tail of the striatum (TOS). Dopamine signaling in the TOS is modulated by internal hunger state through the activity of agouti-related peptide (AgRP) neurons, putative “hunger neurons” in the arcuate nucleus of the hypothalamus. These AgRP neurons are poly-synaptically connected to TOS-projecting dopaminergic neurons through the lateral hypothalamus, the central amygdala, and the periaqueductal grey. We thus delineate a hypothalamic-midbrain circuit that coordinates changes in exploration behavior in the hungry state.

## Introduction

Hunger promotes goal-directed behaviors that are thought to help restore an animal’s homeostatic balance (Smith and Grueter, 2022). Food-seeking is one important behavioral correlate of the hungry state, but there are additional changes in behavior that are not obviously oriented towards obtaining caloric rewards, including decreased anxiety of open spaces, increased tolerance for certain types of pain, increased risk taking in the face of predation, and changes in territorial aggression (Smith and Grueter, 2022). Behavioral adaptation in the hungry state also includes increased exploration and risk taking (Corey, 1978). When an animal is calorically depleted, it will spend more time exploring a maze or novel stimulus (Burnett et al., 2016; Dietrich et al., 2015; Dodt et al., 2024). Increased risk taking and exploration is adaptive to finding potential new food sources and suppressing the aversion of novel stimuli can assist in this foraging.

Although the neural circuits underlying many hunger-induced behavioral changes have been elucidated (Alhadeff et al., 2018; Burnett et al., 2016; Dietrich et al., 2015; Jikomes et al., 2016; Padilla et al., 2016; Salgado et al., 2023), those involved in hunger-induced risk-taking and novel-object exploration are unknown.

Exploration is modulated by the activity of midbrain dopaminergic neurons (DANs) that project to the striatum (Gunaydin et al., 2014; Mikhael and Gershman, 2022). The activity of these midbrain DANs changes when animals learn new information, receive a novel reward, or engage with a novel stimulus (Bromberg-Martin and Hikosaka, 2011; Gunaydin et al., 2014; Krausz et al., 2023; Menegas et al., 2018). Caloric state modulates the magnitude of phasic dopamine (DA) release in the nucleus accumbens (NAcc) and the basolateral amygdala when animals receive overtly rewarding or aversive stimuli, respectively (Alhadeff et al., 2019; Lutas et al., 2019). It is not known, however, whether caloric state modulates DAN responses to a salient, non-caloric stimulus to subsequently change exploratory and novelty-seeking behaviors.

Agouti-related peptide (AgRP) expressing neurons in the arcuate nucleus of the hypothalamus (ARC) modulate food intake and feeding-related behaviors and are activated during fasting (Aponte et al., 2011; Krashes et al., 2011). Here, we show that, consistent with previous studies, hunger increases novel object exploration. The hunger signal that modulates novel object exploration is derived from the activity of AgRP neurons, which, we find, bidirectionally modulate DA signaling in the tail of the striatum (TOS), a nigrostriatal DA signaling pathway thought to represent salience and threat prediction, during exploration (Akiti et al., 2022; Green et al., 2024; Menegas et al., 2018). We provide causal evidence that this change in exploration in the hungry state is due to changes in TOS DA release. Thus, we describe a hypothalamic-midbrain neuronal circuit that concomitantly regulates exploration with caloric need.

## Results

### Hunger restructures spontaneous open field exploration behavior

To study how hunger modulates exploration, we used a two-stage, open-field novel object exploration (NOE) assay in which mice were first either food-restricted (hungry mice) or given food *ad libitum* (sated mice) (see Methods) and allowed to freely explore an empty arena for two days (H1 and H2) before an object is placed in one section of the arena (Figure 1A) (Akiti et al., 2022). Mouse movements and posture were captured with an overhead camera and analyzed with Motion Sequencing (MoSeq), an unsupervised machine-learning based algorithm that segments continuous animal behavior into characteristic, discrete “syllables” (Figure 1B) (Lin et al., 2024; Wiltschko et al., 2015). Hungry and sated animals employ behavioral syllables at different frequencies in their spontaneous behavior in both types of sessions (Figure 1C), consistent with state-dependent changes in behavior (Hochbaum et al., 2024; Markowitz et al., 2018).

**Figure 1:**
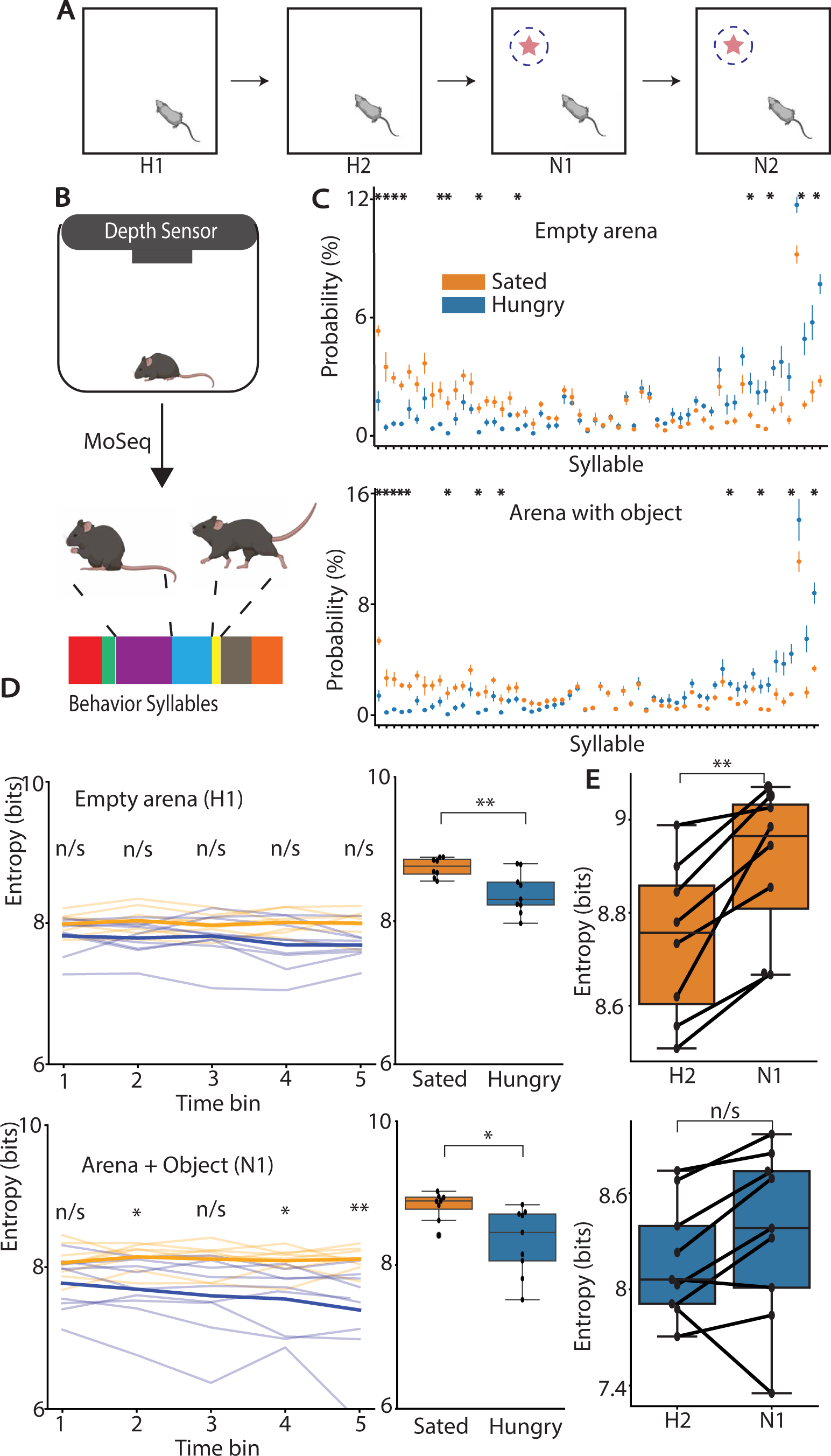
Hunger restructures spontaneous exploration behavior. **A)** Schematic representation of day-to-day setup for open-field novel object exploration assay. Star indicates the novel object, dashed blue lines indicate interacting radius. **B)** MoSeq workflow to capture animal behavioral data and segment it into individual syllables. **C)** Changes in individual syllable frequency between hungry and sated animals when animal explores an empty arena (days H1-H2, top) and when it explores an arena with an object (days N1-N2, bottom). n=9 hungry animals, n=8 sated animals, two sessions/animal, *:p<.05, Mann-Whitney U-test with Bonferroni correction.) Values are normalized to total syllable usage across all animals in each type of session. **D)** Syllable transition entropy for animals during the first session without object present divided into quintiles (top left; n/s: p>.05, Mann-Whitney U-test with Bonferroni correction) and compared across groups (top right) averaged across sessions H1-2, and similar comparisons with the object present (bottom row; Mann-Whitney U-test with Bonferroni correction). **E)** Changes in syllable transition entropy on second day of exploring an arena to first day of exploring arena with object in sated (top) and hungry (bottom) animals.

MoSeq describes continuous behavior as a Markovian process, allowing for analysis of the transitions between syllables and of the statistics of the distribution of syllable transitions. In particular, the entropy of the syllable transition distribution (syllable transition entropy) measures the randomness of an animal’s behavior, such that lower entropy corresponds to more predictable behavior (Wiltschko et al., 2015). The syllable transition entropy of hungry animals is lower than that of sated animals both in the presence (U=64, p=0.0055) and absence of the object (U=62, p=0.011) (Figure 1D).

Thus, hungry mice exhibit less variability in the sequence of syllables used during spontaneous movements. Additionally, the presence of the object decreases the behavioral structure of sated (Wilcoxon paired signed-ranked test; W=0, p=0.008), but not hungry (W=10, p=0.164), animals (Figure 1E), indicating that hunger alters the impact that a novel object has on spontaneous behavior in mice.

### Hunger changes exploration of a novel object in a familiar arena

Mice explore novel objects using characteristic approach-avoid bouts (Menegas et al., 2018) which can be subdivided into those in which the animal keeps its tail behind its nose the entire time (tail behind bouts) or those in which the animal exposes its tail to the object (tail exposed bouts) (Figure 2A-B). These two modes of interaction are identified based on video key-point tracking and their differential expression reflect changes in risk assessment behavior (Akiti et al., 2022; Mathis et al., 2018). We find that hungry animals (n=9) spend more time exploring a novel object than sated animals (n=8) (Interacting Time: U=63, p=0.008), and while interacting with the object, spend a greater fraction of that time with their tail exposed to the object (normalized Tail Exposure Time: U=71, p=0.0009), but do not approach the object more often (Number Interactions: U=47, p=0.31) (Figure 2A,C). Furthermore, each animal’s syllable transition entropy on the first day of novel object exploration (day N1) is inversely correlated with some, but not all, metrics of exploration of the object (Figure 2D). This relationship suggests a potential common neural process mediating differences in behavioral structure and differences in novel object exploration between hungry and sated animals.

**Figure 2:**
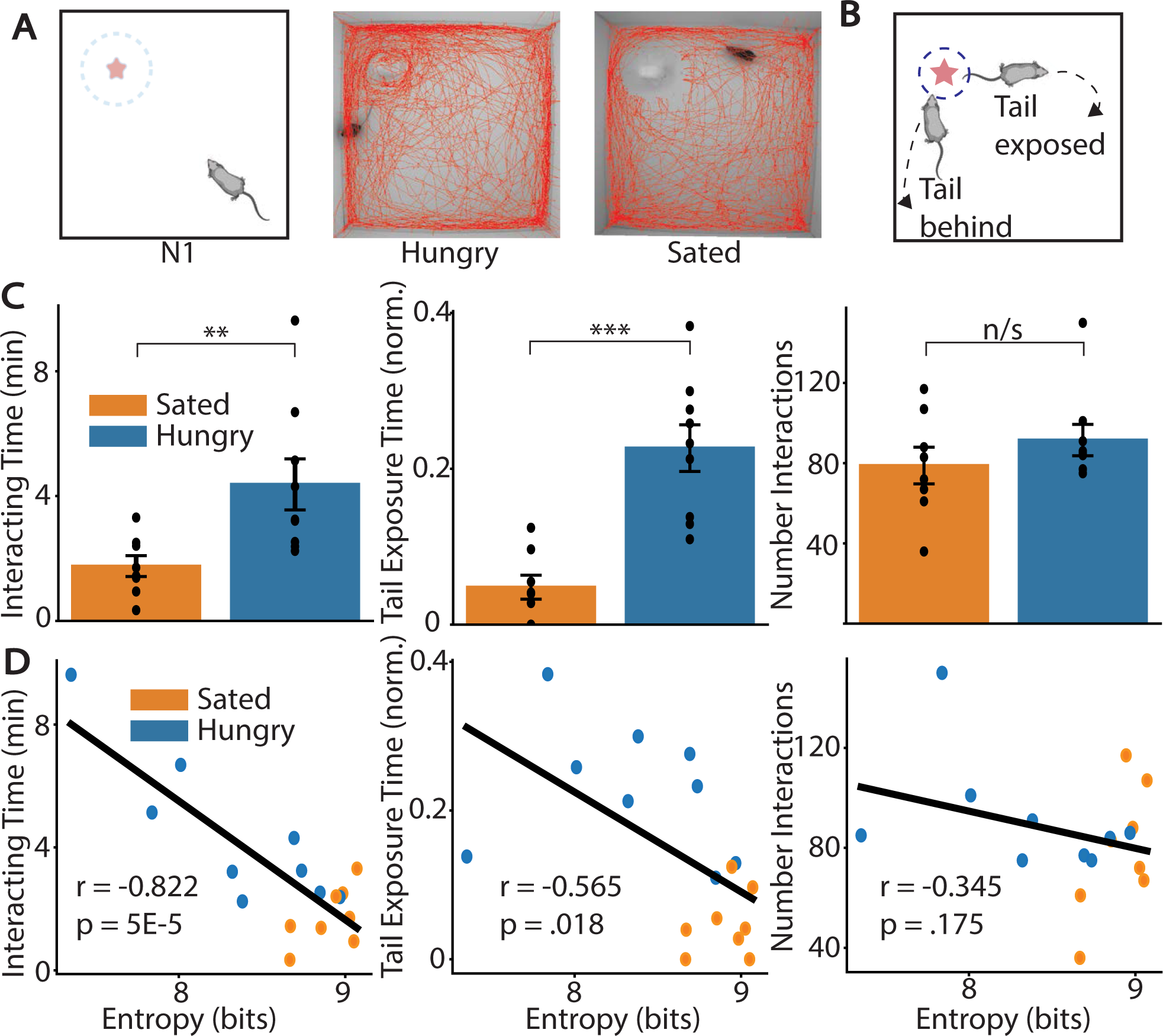
Hunger increases directed interaction and reduces risk assessment of a novel object. **A)** Schematic of animal exploration of arena with a novel object present in one corner of the arena (left). Comparison of the position (red trace) of a hungry (middle) and sated (right) animal on the first day of exposure to the novel object (N1). **B)** Mice retreat from a novel object either with tail behind the entire time (bottom left) or with their tail exposed (top right). **C)** Comparison of three different novel object exploration statistics – left: total time animal spends near the object (Interacting Time), middle: the fraction of the time the animal is around the object that their tail is closer than their nose (Tail Exposure Time norm.), and right: the number of times the animal interacts with the object (Number Interactions). **D)** Regression of the novel object exploration metric (left: Interacting time, middle: Tail exposure time normalized, right: Number of interactions) and syllable transition entropy across animals on day N1. Pearson’s correlation coefficient was used.

### Hunger modulates dopamine transients in the TOS during novel object exploration

The mesostriatal dopaminergic system regulates novel object exploration behavior and changes in behavioral structure (Akiti et al., 2022; Gunaydin et al., 2014; Markowitz et al., 2023). In particular, DA is released in the TOS when mice retreat from a novel object. The magnitude of this increase is inversely correlated with the amount the animal explores a novel object, leading to the hypothesis that DA release in the TOS reflects a threat-prediction error, crucial for learning about potential threats in the environment (Akiti et al., 2022). We hypothesized that increased novel object exploration in hungry animals is caused by reduced phasic DA signaling in the TOS. To test this hypothesis, we used adeno-associated virus (AAV) to express a new genetically encoded sensor for DA, dLight3.8 (Roshgadol et al., in preparation), in the TOS and implanted an optic fiber to record changes in fluorescence that are indicative of DA release (Figure 3A, S2C). Upon ligand binding dLight3.8 exhibits robust increases in fluorescence intensity and lifetime (Lodder et al., in preparation; Roshgadol et al., in preparation). Fluorescence lifetime is insensitive to many artifacts that plague intensity measurements such as differences in sensor concentration, fiber placement, photobleaching and hemodynamics (Lodder et al., in preparation; Lee et al., 2021; Simpson et al., 2024). Therefore, we recorded dLight3.8 fluorescence lifetime as animals explored the novel object as a proxy for fluctuations in DA concentration in the TOS using **f**luorescence **li**fetime **p**hotometry (FLiP) (Lodder et al., in preparation).

**Figure 3:**
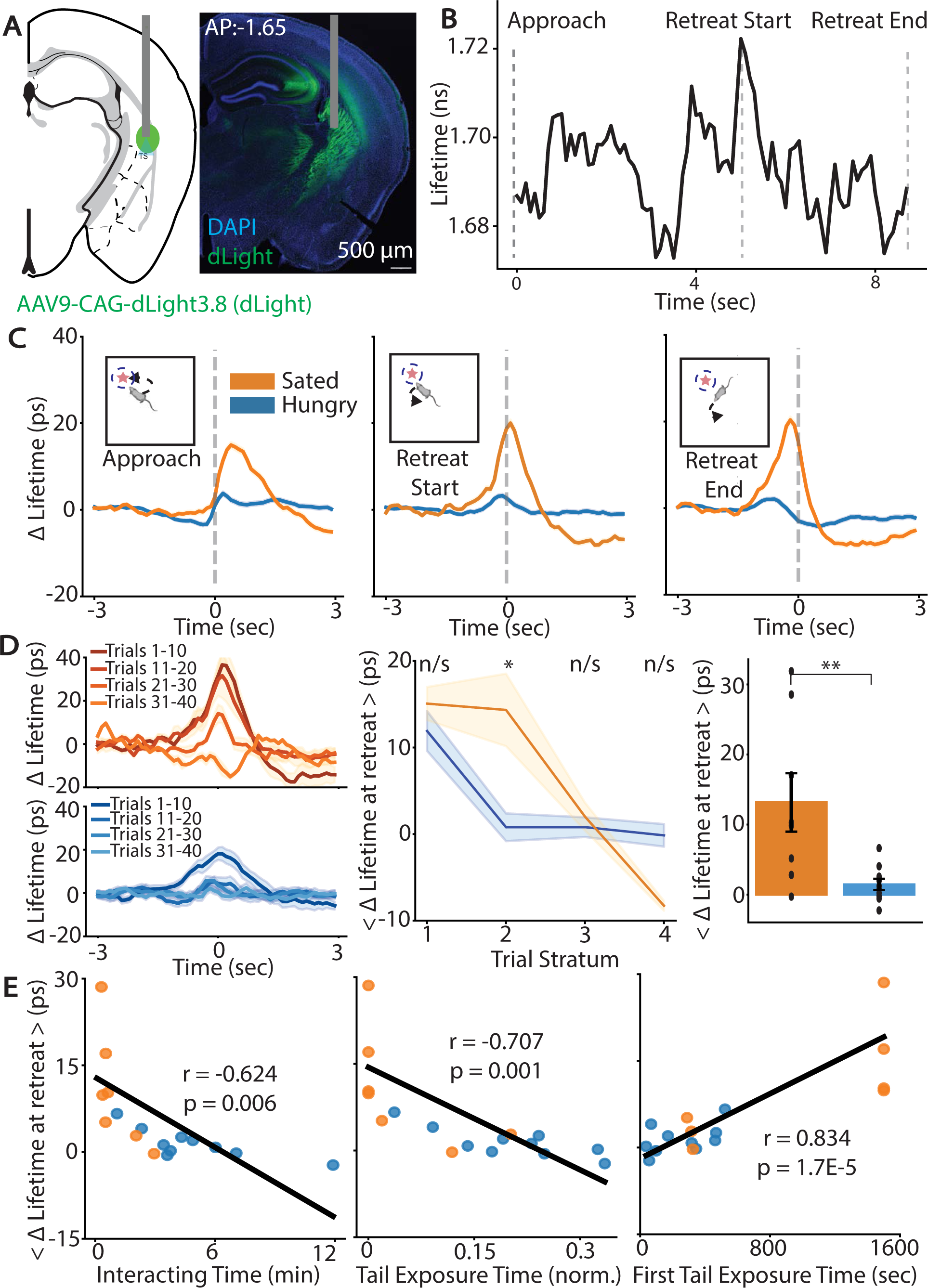
Hunger decreases dopamine release in the tail of the striatum during novel object assessment. **A)** Experimental design schematic and representative image for injection/surgery in the tail of the striatum (TOS) (left) and an example histology of fiber placement (right). Gray bar indicates fiber optic placement. **B)** dLight3.8 fluorescence lifetime fluctuations in the TOS when animals interact with a novel object as recorded using FLiP. **C)** Hunger modulation of dLight3.8 fluorescence lifetime fluctuations when animals interact with a novel object. Graphs are aligned to different stages of interaction with a novel object when an animal approaches the object (Approach), when an animal begins to move away from the object (Start Retreat), and when an animal stops interacting with the object (End Retreat). Bold line represents the mean and shaded area is the standard error around the mean (SEM) calculated across animals and bouts for all photometry graphs unless otherwise specified. **D)** Difference in dLight3.8 fluorescence lifetime fluctuations as animals make repeated interactions with the object (“trials”). The dLight3.8 signal aligned to start of retreat, grouped into strata of 10 trials across sated and hungry animals (left; bold line is mean, shaded area is SEM calculated across animals), the change in mean lifetime signal around the start of retreat across these trial strata (middle; Mann-Whitney U-test with Bonferroni correction, *:p<0.05), and the overall difference in retreat-associated lifetime signal between hungry and sated animals (right). **E)** Regression of the mean dLight3.8 fluorescence lifetime signal during retreat for an animal with its exploration of a novel object. Pearson’s correlation coefficient was used.

Exploration of the object modulates the TOS dLight3.8 fluorescence lifetime signal, such that the fluorescence lifetime rises when an animal approaches the novel object, peaks when the animal begins to retreat from the object and decreases as the animal fully retreats from the object (Figure 3B).

To assess if hunger alters TOS DA release, we measured dLight3.8 fluorescence lifetime in the TOS on the first day of novel object exploration (N1) in the NOE assay. dLight3.8 fluorescence lifetime transiently increases upon retreat from the novel object in both hungry and sated mice, though the magnitude of this signal is lower in hungry animals compared to sated animals (Figure 3C). In sated, but not hungry, animals, the fluorescence lifetime signal transiently dips below the baseline shortly after the start of retreat (Figure 3C). This dip is present predominantly in the lifetime signal during the first several interactions with the object (Figure 3D). The transient increase in fluorescence lifetime signal around the object decreases over subsequent novel object exploration bouts in both hungry and sated animals (Figure 3D), consistent with habituation to the presence of the object. The magnitude of the retreat-associated fluorescence lifetime signal on average (<Δ Lifetime at retreat>) is comparable between hungry and sated animals in the first several interactions but decays more rapidly over bouts of object exploration in hungry animals compared to sated animals (Figure 3D). Over the entire session, hungry animals compared to sated animals have, on average, smaller fluorescence lifetime transients (reflecting decreased DA release) in the TOS upon retreat from the novel object (U=11, p=0.008, n=10 hungry, n=8 sated animals) (Figure 3D). There is no difference in lifetime signals between hungry and sated animals for a ligand-binding mutant version of the dLight3.8; thus, the differences in the DA signal between the groups of animals are not driven by ligand-independent effects on fluorescence lifetime (U=6, p=0.7; n=3 sated, n=3 hungry animals) (Figure S2A).

Furthermore, when restricting analysis to interactions in which both groups of animals did not expose their tail at all, hungry animals still displayed a lower TOS DA signal compared to sated animals (U=11, p=0.008) (Figure S2B), indicating that differences in the DA signal are not driven by gross behavioral differences between the two groups of animals.

We observed significant variation in the magnitude of the fluorescence lifetime signal in hungry and sated animals exploring the novel object. The variation in the magnitude of the fluorescence lifetime signal when an animal retreats from the novel object is correlated with that animal’s exploration of the object, and this correlation is preserved in both hungry and sated animals (Figure 3E). These results suggest that changes in novel object exploration behavior in hungry animals are due to modulation of TOS DA signaling.

### AgRP neurons bidirectionally modulate exploration behavior and TOS DA transients

AgRP-expressing neurons in the ARC are activated by fasting and increased activity of these neurons decreases anxiety and increases risk-taking behavior (Dietrich et al., 2015; Dodt et al., 2024; Liu et al., 2023; Mandelblat-Cerf et al., 2015; Padilla et al., 2016). AgRP neurons regulate DA release in the NAcc via a polysynaptic projection through the lateral hypothalamus (Alhadeff et al., 2019; Grove et al., 2022). We hypothesized that AgRP neurons are the neural source of the hunger drive that modulates exploration and TOS DA signaling. The activity of AgRP neurons can be manipulated with designer receptors exclusively activated by designer drugs (DREADDs) in AgRP-*ires*-Cre mice (Krashes et al., 2011; Tong et al., 2008). Thus, to test the role of AgRP neural activity in hunger modulation of exploration and TOS DA signaling, we either stimulated or inhibited AgRP neurons using DREADDs in sated or hungry mice, respectively, and recorded resulting changes in exploration and TOS DA signaling. To simultaneously manipulate AgRP neurons and record DA signaling, we utilized AgRP-*ires*-Cre mice injected with (1) a Cre-dependent AAV in the ARC to express either excitatory (hM3Dq) or inhibitory (hM4Di) DREADDs in AgRP neurons and (2) a Cre-independent AAV in the TOS to express the dLight3.8 sensor. We also implanted a fiber optic in the TOS to perform frequency-modulated intensity fiber photometry to measure DA fluctuations in the TOS (Figure 4A).

**Figure 4:**
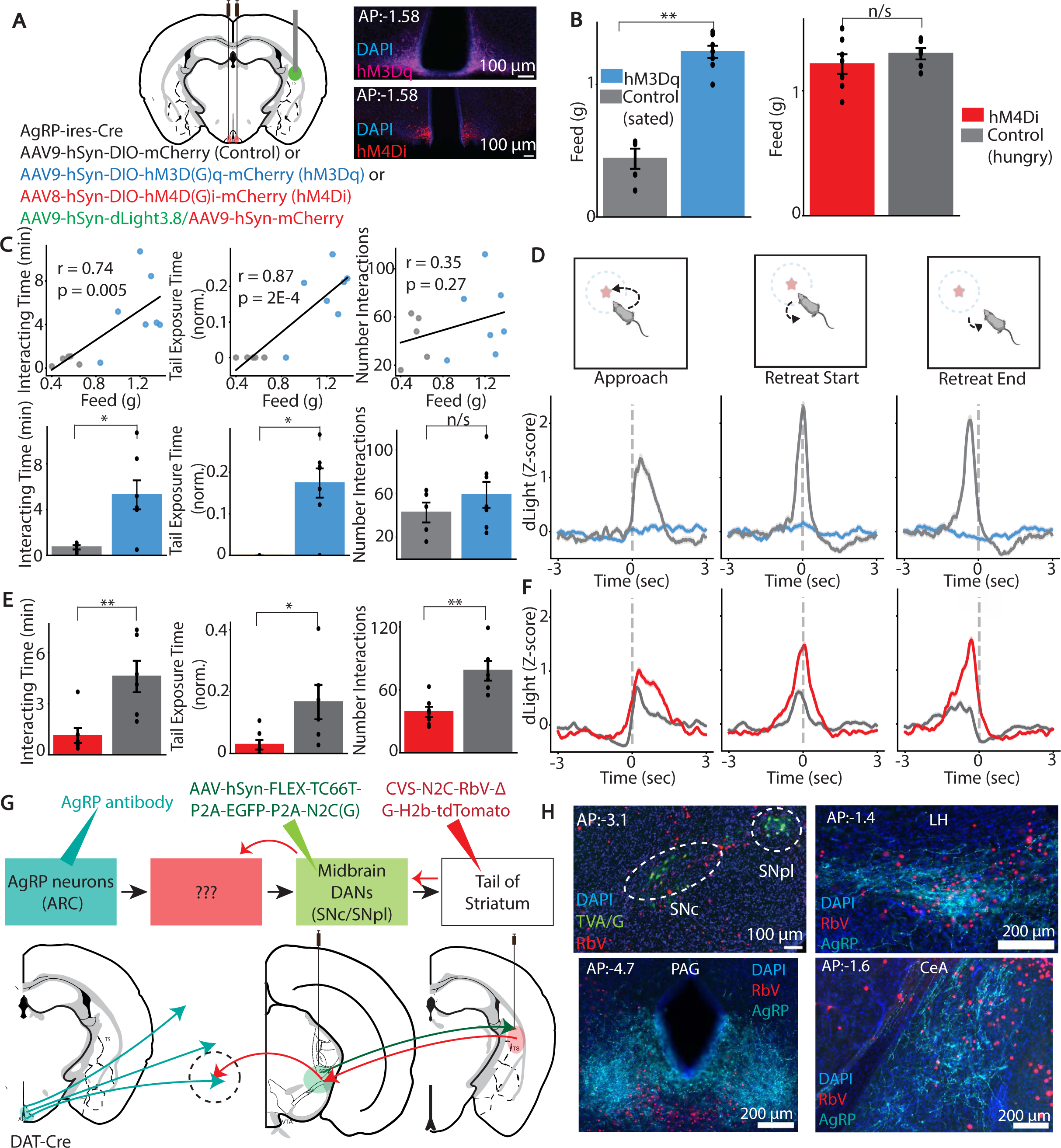
AgRP neurons modulate novelty exploration and risk assessment and associated DA fluctuations in the TOS. **A)** Injection scheme and representative image for recording DA in the TOS and manipulating AgRP neurons with either excitatory or inhibitory DREADDs (left). **B)** Effect of hm3Dq-mediated AgRP neuronal excitation in sated animals on feeding behavior (left) and effect of hm4Di-mediated AgRP neuronal inhibition in hungry animals on feeding behavior (right). **C)** Regression of effect of excitation of AgRP neurons in sated animals on feeding from B and exploration/risk assessment of a novel object (top; Pearson’s correlation coefficient was used). Effect of excitation of AgRP neurons on exploration/risk assessment of a novel object (left: Interacting time; middle: Tail Exposure Time norm.; right: Number Interactions) (bottom). **D)** Effect of excitation of AgRP neurons in sated animals on dLight3.8 fluorescence intensity fluctuations in the TOS during interactions with a novel object. **E)** Effect of inhibition of AgRP neurons in hungry animals on exploration and risk assessment of a novel object. **F)** Effect of inhibition of AgRP neurons in hungry animals on dLight3.8 fluorescence intensity fluctuations in the TOS during interactions with a novel object. **G)** Schematic of rabies tracing experiment to find putative links between AgRP neurons and TOS-projecting DANs (top) with injection schema overlaid over coronal section schematics (bottom) **H)** Starter population of DANs in the SNpl that are double positive for both TVA/G-protein (green) and RbV (red) (top left) and regions of overlap between AgRP neuron axons (cyan) and RbV+ neurons (red) – the LH (top row; right), the PAG (bottom row; left), and the CeA (bottom row; right).

We hypothesized that stimulation of AgRP neurons in sated mice would both increase an animal’s feeding behavior and increase its exploration of a novel object.

Chemogenetic activation of AgRP neurons in sated mice robustly increases feeding on calorically rewarding substances (U=35, p=0.002; n=5 control sated, n=7 hM3Dq experimental animals) (Figure 4B). Additionally, in a separately tested behavioral context, chemogenetic activation of AgRP neurons in sated mice also increases the amount of time an animal explores the novel object (U=32, p=0.018), and the fraction of time it spends with its tail exposed towards the object (U=32.5, p=0.012) (Figure 4C).

AgRP neuron activation does not change the number of times an animal approaches the object (U=22.5, p=0.464) (Figure 4C), similar to the behavioral phenotype in calorically depleted mice (Figure 2C). Notably, the effect sizes of the chemogenetically-induced feeding and chemogenetically-induced change in exploration are correlated, animal-by-animal (Figure 4C), consistent with variance in transfection of AgRP neurons with the stimulatory DREADD influencing both food intake (Aponte et al., 2011; Betley et al., 2015) and exploration behavior. AgRP neuron stimulation also decreased the amplitude of dLight3.8 fluctuations in the TOS when animals retreated from the novel object (Figure 4D), similar to caloric depletion (Figure 3C, D).

Conversely, inhibition of AgRP neurons in hungry animals decreases exploration of the novel object (U=40, p=0.004; n=6 control hungry, n=7 hM4Di experimental animals) and tail exposure towards the object (U=38, p=0.018) (Figure 4E). Inhibition of AgRP neurons in hungry animals decreases the number of interactions an animal makes with the object (U=40, p=0.004) (Figure 4E), though modulation of AgRP neural activity does not change average speed in the arena (hM3Dq: U=26, p=0.202; hM4Di: U=21, p=1) (Figure S3B,D). Furthermore, the decreased exploration is accompanied by increased TOS DA signaling upon retreat from the novel object, (Figure 4F), although the increase in TOS DA signaling by AgRP inhibition is more modest than the reduction induced by AgRP excitation (Figure 4D). The decrease in novel object exploration and increase in TOS DA signaling is not accompanied by changes in feeding behavior with AgRP neural inhibition (U=22, p=0.445) (Figure 4B). Given that the activity of only a small percentage of AgRP neurons is necessary to coordinate feeding behavior, the lack of change in feeding behavior is likely due to incomplete transduction of all AgRP neurons (Aponte et al., 2011). The segregation of effects on feeding and exploration suggests that the observed changes in exploration are not a simple byproduct of the feeding drive.

To uncover the neuronal circuitry by which AgRP neurons may modulate TOS-projecting DANs, we performed projection-specific rabies virus tracing of monosynaptic inputs to TOS-projecting DANs. For this experiment, we used animals that express Cre recombinase under control of the dopamine transporter (DAT) promoter (DAT-*ires*-Cre). Cre expression in this mouse is highly selective for and penetrant in DANs of the substantia nigra and ventral tegmental area (VTA) (Bäckman et al., 2006). We injected an AAV carrying a Cre-dependent bicistronic construct encoding both a weakened EnvA receptor, TVA TC66T (Miyamichi et al., 2013) and G protein in the substantia nigra of DAT-*ires*-Cre mice, causing expression of TVA and G proteins in midbrain DANs. Four weeks later, we injected animals with EnvA-pseudotyped and G-deleted rabies virus (RbV) of the lower-toxicity CVS-N2C strain (CVS-N2C-RbV-ΔG-H2b-tdTomato) in the TOS (Reardon et al., 2016). The RbV(EnvA) specifically infects DAN axons in the TOS that express the TVA TC66T, and once in the DANs, RbV are amplified in DANs that have been complemented by G, and transmitted to neurons presynaptic to TOS-projecting DANs (Figure 4G). After waiting six days, we sacrificed animals and immunohistochemically stained for AgRP and examined the colocalization of AgRP+ axons and RbV+ neurons across brain regions (Figure 4G). As expected, primary infection (RbV+/TVA+) is restricted mostly to DANs in the substantia nigra pars lateralis (SNpl) (Figure 4H, S3E). There are three primary sites of colocalization between AgRP+ axons and RbV+ neurons: the lateral hypothalamus (LH), the central amygdala (CeA), and the periaqueductal grey (PAG) (Figure 4H). In contrast, there is minimal colocalization in other major projection sites of AgRP neurons (Figure S3F). In the absence of G protein, the pseudotyped RbV infects DAN axons in the striatum and moves to DAN cell bodies but did not travel to the abovementioned regions, confirming the necessity of G-complementation (Figure S3G). This anatomical tracing delineates neural pathways through which AgRP neurons may influence the activity of TOS-projecting DANs to change an animal’s exploration behavior.

### TOS DA transients modulate exploration in hungry animals

To determine if the hunger-induced changes in DA signaling in the TOS produce the change in exploration behavior, we examined if optogenetically stimulating DA release in the TOS during interactions with a novel object decreases an animal’s exploration behavior. To target TOS-projecting DANs, we employed a retrograde strategy using a serotype of AAV, AAVDJ/9, which efficiently infects DAN axons (Figure S4A; Düring et al., 2020). In DAT-*ires*-Cre mice, we bilaterally co-injected two viruses in the TOS: (1) an AAVDJ/9 to express Cre-dependent ChRmine (a sensitive red-shifted opsin; Marshel et al., 2019) in DANs and (2) an AAV to express Cre-independent dLight3.8 sensor in TOS. In control animals, the Cre-dependent ChRmine construct was replaced by a non-fluorescent control protein (10xmyc) (Figure 5A). We implanted fiber optics in both the TOS and over the SNpl (the primary site from which TOS-projecting DANs originate) (Figure 4H, 5A; Menegas et al., 2018). This approach permits simultaneous stimulation of DAN cell bodies via the SNpl fiber and measurement of resulting DA release in the TOS via the TOS fiber.

**Figure 5:**
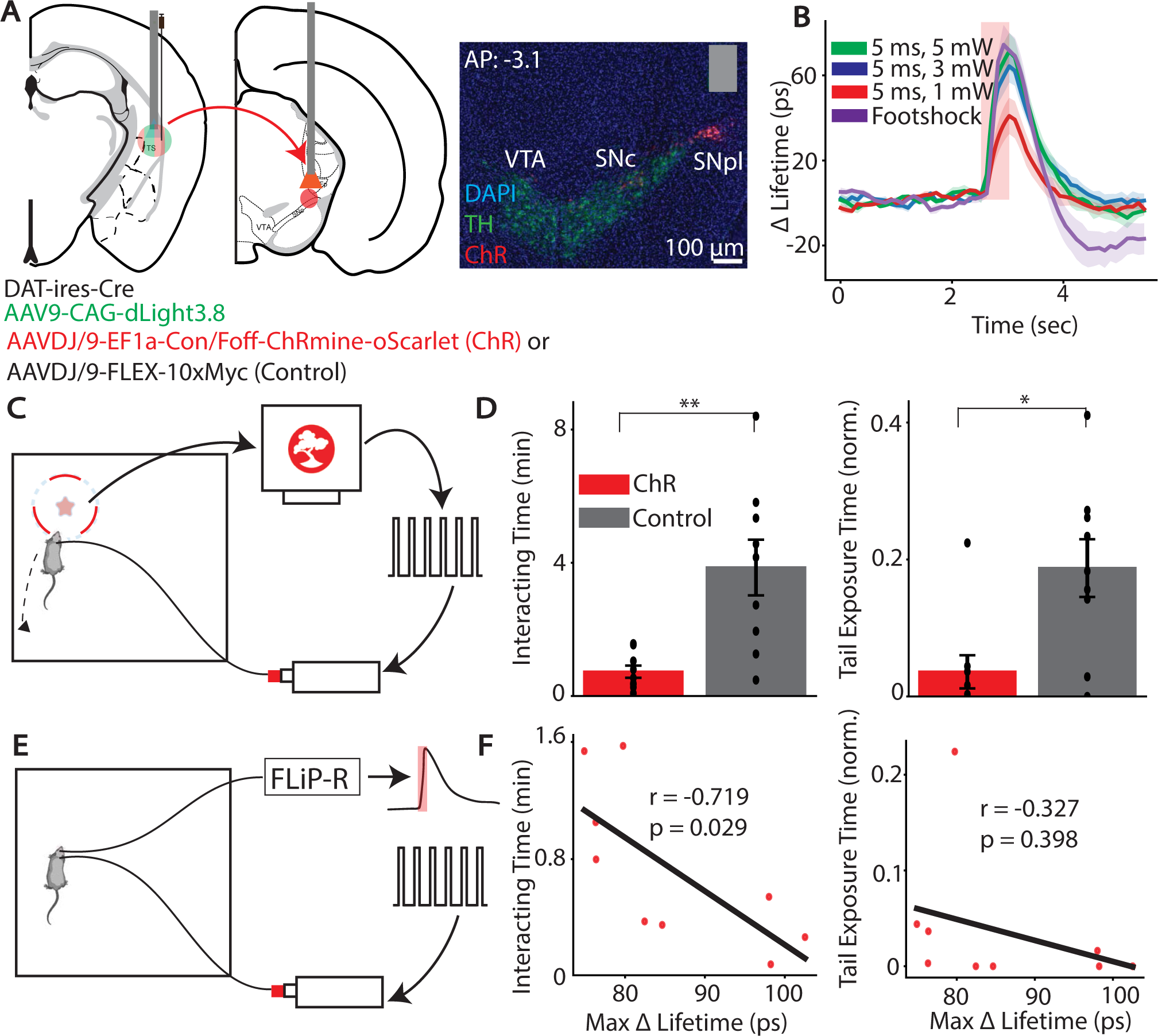
TOS DA modulates novelty exploration and risk assessment behavior in hungry animals. **A)** Injection and surgery schematic for manipulating TOS-projecting dopaminergic neurons and recording and resulting release (left) and example histology placement of stimulating fiber in the midbrain (right). Gray bar indicates fiber optic placement. **B)** Calibration of optogenetic stimulation parameters to evoke similar dopamine release as a footshock. Red bar represents time of either optogenetic stimulation or footshock. **C)** Schematic of closed-loop optogenetic activation of TOS-projecting DANs when animal approaches a novel object (star). The animal’s presence within a given ROI around the object (red dashed circle) is processed in real-time using the Bonsai-rx software (top computer), which then generates a pulse train (right) to allow for pulsed activation of a laser (bottom), which is transmitted to the mouse. **D)** Effect of optogenetic activation of TOS-projecting DANs in hungry mice on novel object exploration behavior. **E)** Schematic of post-hoc validation of evoked DA release in the TOS with optogenetic stimulation using FLiP in a separate open field arena without the object present. The same optogenetic pulse pattern as was used in the behavioral assay was fed into a laser (right pulse pattern and laser). Simultaneously, on the ipsilateral hemisphere, fluorescence lifetime signal was collected from the fiber implanted in the TOS to record fluorescence lifetime fluctuations in the TOS (left mouse). The signal was generated by the FLiP system (top) and analyzed post-hoc (right trace with overlying red rectangle representing the time of optogenetic activation.) **F)** Regression of change in exploration and risk assessment with level of DA release evoked by the optogenetic stimulation parameter across mice. Pearson’s correlation coefficient was used.

We calibrated the optogenetic stimulation to mimic the level of TOS DA release evoked by a natural stimulus. Since DA release in the TOS is thought to reflect a threat-prediction error, we exposed an initial cohort of hungry animals to a footshock, a robustly threatening stimulus, to drive DA release in the TOS (Figure 5B). We then stimulated TOS-projecting DANs in mice expressing ChRmine at the SNpl and recorded resulting TOS DA release. We varied the optogenetic stimulation light power, frequency, and pulse width to determine a set of optogenetic stimulation parameters that evoked DA release in the TOS of similar amplitude to the footshock (Figure 5B; n=4 sites across n=3 animals).

To perform closed-loop manipulation of TOS DA signaling during behavior, hungry animals were allowed to explore the novel object while their position was tracked using a real-time behavioral monitoring system, Bonsai-rx (Lopes et al., 2015). When a portion of the animal’s body fell within 7 cm of the object, the optogenetic stimulation was delivered (Figure 5C). ChRmine expressing animals (n=9) under closed loop optogenetic activation of DANs spend less time interacting with the object (U=74, p=0.004) and a smaller fraction of that time with their tail exposed towards the object (U=58, p=0.016) than control hungry animals (n=9) experiencing the same stimulation paradigm (Figure 5D). This reduction in exploration is not driven by alteration of the food-seeking drive in opsin expressing mice as these mice eat the same amount as control hungry mice in a feeding test, conducted immediately after the NOE assay (U=5, p=0.689; n=9 control hungry animals, n=8 opsin experimental animals) (Figure S4C).

To understand whether variance in exploration in the opsin-expressing mice could derive from variance in the evoked TOS DA signal, we used FLiP to measure optogenetically-evoked DA release for each animal in a different open arena without the object present (Figure 5E). The level of evoked TOS DA release in the open field inversely correlates with time spent exploring the object (Figure 5F), suggesting that the TOS DA signal can modulate exploration behavior. In contrast, although optogenetic activation of TOS-projecting DANs decreases tail exposure, the variation in the amount of tail exposure was not correlated with the amount of DA released (Figure 5F).

Together, our data support that TOS DA release reflects threat-prediction, and that stimulating this system is sufficient to cause hungry animals to avoid the novel object.

## Discussion

In this study, we describe a neural circuit underlying changes in exploration behavior in hungry animals. Although it is intuitive that hunger causes animals to explore their environment and take greater risks in order to find food, the neural circuits mediating this behavioral change had not been elucidated. We find that caloric state modulates phasic DA signals in the TOS evoked when animals interact with a novel, non-caloric object. This effect results from the influence of AgRP neurons in the ARC over TOS DA signaling. Furthermore, bidirectional manipulations of the activity of AgRP neurons alter both novel object exploration and TOS DA signaling. This hunger-induced suppression in TOS DA signaling is causal for the increased exploration behavior in hungry animals. We thus delineate a circuit that coordinates exploration and risk-taking behavior with caloric need.

### Hunger modulation of spontaneous and directed exploratory behavior

Internal states such as hunger change animal behavior; however, these changes have primarily been studied in environments where food was once present (e.g., an animal’s home environment) or in the presence of food itself (Burnett et al., 2016; Dietrich et al., 2015; Krashes et al., 2011; Liu et al., 2023). Instead, we investigated how hunger changes behavior in the absence of overt food cues. We find that hunger increases the structure of spontaneous behavior, increases directed exploration and decreases risk assessment of a salient, novel object. These behavioral metrics are correlated, suggesting a common neural process controlling both object exploration and behavioral structure. The more structured behavior, as reported by MoSeq syllable transition entropy, in hungry animals is consistent with increased spatial exploration using a strategy that efficiently explores physical space with repetitive, stereotyped movements that limitedly explore behavioral space (e.g., spiral search) (Müller and Wehner, 1994). In contrast to prior results (Krashes et al., 2011) we do not observe hunger-evoked changes in average speed, potentially due to the lack of food-signaling cues in our experiments.

### Hunger modulation of a nigrostriatal dopamine signaling pathway through AgRP neurons

DA transients in the TOS signal threat-prediction errors (in contrast to those in the ventral striatum which may signal reward prediction errors; Menegas et al., 2017), such that lesions of TOS-projecting DANs decrease an animal’s risk assessment and allow them to more quickly overcome threats in pursuit of perceived reward (Akiti et al., 2022; Menegas et al., 2018; Tsutsui-Kimura et al., 2022). This DA response integrates features of the salient stimulus that evoke the DA release, such as size of threatening object or magnitude of tone (Menegas et al., 2018; Tsutsui-Kimura et al., 2022). We find that this DA response is also modulated by internal state, such that hungry animals have decreased DA release in the TOS when interacting with a novel object, as measured using fluorescence lifetime photometry. Under the threat-prediction error framework, we propose that hungry animals explore a novel stimulus more because the DA release in the TOS is lower, allowing the animals to learn more quickly that the novel stimulus, a potential threat, will not cause them harm. Thus, hunger facilitates exploration in the face of potential threat by suppressing DA dependent threat estimates in the TOS. Similar fasting-induced decreases in DA release occur in the basolateral amygdala in response to an aversive tail shock in mice (Lutas et al., 2019).

Hunger increases the magnitude of DA response in the NAcc to food through the activity of AgRP neurons, which disynaptically project to VTA DANs through the LH (Alhadeff et al., 2019; Cone et al., 2014; Grove et al., 2022). Activation of AgRP neurons increases feeding and exploration of a novel object; notably we find these two measures to be correlated animal-by-animal, suggesting that AgRP neural activity is causal for the changes in exploration in hunger. This causality is further established by bidirectional manipulation of AgRP neurons which changes both novel object exploration and associated TOS DA signaling. The AgRP-induced suppression of TOS DA release in the NOE assay is opposite from the AgRP-induced potentiation of DA release in the NAcc in response to food, indicating that internal state modulation of DA signaling is dependent on the nature of the stimulus evoking the DA release (Alhadeff et al., 2019). Notably, inhibition of AgRP neurons has a more subtle effect on exploration and TOS DA signaling than activation of these neurons, potentially due to incomplete inactivation of AgRP neurons and the ability of a small fraction of AgRP neurons to coordinate aspects of hunger and feeding (Aponte et al., 2011). Chemogenetic inhibition of AgRP neurons similarly has a subtle effect on modulating the activity of downstream corticotropin-releasing hormone neurons in the paraventricular hypothalamus (Douglass et al., 2023). AgRP neurons disynaptically project to TOS-projecting DANs through the LH, CeA, and PAG, though it is possible that AgRP neurons regulate the activity of TOS-projecting DANs through a different polysynaptic pathway (Livneh et al., 2017), or even through neuropeptide release into the ventricles. These possibilities present promising directions for future studies.

### Dopaminergic signaling in the TOS changes exploration behavior in hunger

Hunger modulates several neural systems involved in anxiety and pain processing (Alhadeff et al., 2018; Padilla et al., 2016). Moreover, DA signaling in the TOS assists in learning from some, but not all, aversive and salient stimuli (Menegas et al., 2018).

Thus, there are multiple possible neural circuits that may coordinate changes in novel object exploration in hungry animals other than DA signaling in the TOS. Nevertheless, closed-loop, calibrated optogenetic stimulation of TOS-projecting DANs in hungry animals when they explore a novel object is sufficient to reduce novel object exploration. Importantly, the variance in exploration behavior was correlated with the level of optogenetically-evoked TOS DA release. Coupled with our finding that hunger suppresses the TOS DA response at novel object exploration, we propose that changes in novel object exploration behavior in hungry animals are due to modulation of TOS DA signaling.

Previous studies optogenetically manipulated DA release in the TOS but did not calibrate the magnitude of the DA release to match a behaviorally evoked signal (Greenstreet et al., 2022; Menegas et al., 2018; Schmack et al., 2021; Zafiri et al., 2023). We combined the AAVDJ/9 serotype to retrogradely specifically target TOS-projecting DANs and used absolute measurements of DA using FLiP to calibrate the optogenetic stimulation parameter to mimic DA release in the TOS evoked by a footshock. This calibration was necessary as supraphysiological levels of evoked DA release may cause different behavioral effects from physiological levels of DA release (Coddington et al., 2023; Long et al., 2024; Millard et al., 2024).

In this study, we identify a hypothalamic-midbrain circuit that underlies the long-known ethological adaptation of increased exploration in the hungry state. This work sheds light on how dopaminergic circuitry integrates internal state to change how an animal learns to overcome threats and change its exploration.

### Methods Animals

Male and female mice aged 9-20 weeks were used. The following mouse lines were used: C57BL6/J (The Jackson Laboratory, 000664); *AgRP-ires-Cre* (The Jackson Laboratory, 012899); *DAT-ires-Cre* (The Jackson Laboratory, 006660). All transgenic mice were used as heterozygous and bred inhouse on a congenic C57BL/6J background. We did not observe any significant differences in behavior between the sexes and all experiments contained animals of both sexes. Animals were group-housed until the day before experimentation began on a 12-hour reversed light/dark cycle with standard chow and water provided *ad libitum*. During behavioral experimentation, animals were single housed and for hungry animals, animals were food restricted to 80-90% of their *ad libitum* baseline weight. Behavioral tests were conducted between 12:00 and 19:00 each day, and animal behavior was assessed at roughly the same time day-by-day. All animal care and experimental procedures were approved by the Harvard Standing Committee on Animal Care following guidelines described in the US NIH Guide for the Care and Use of Laboratory Animals.

### Novel Object Exploration and Refeeding Assay

An open-field, freely-moving novel object exploration behavioral assay was implemented as described previously (Akiti et al., 2022). A flat open field arena ∼60 cm x 60 cm was used for experimentation. For behavioral experiments using Motion Sequencing, a black arena was lit with an overhead white light to visualize black mice on a black background. For all other experiments, the arena was white and illuminated with infrared LED strips. Briefly, the animals were habituated to handling for at least three days for 30 minutes per day. During handling, animals without implants or without injections were scooped using a transfer box. A transfer box was put into the corner of the animals’ cage and animals were allowed to freely enter it. Once the animal entered the box, the box was tilted up and replaced back on the floor of the cage. Handling ended once either 30 minutes had passed or the animal had entered the box five times (Akiti et al., 2022). If animals had a fiber optic implant, handling instead consisted of briefly restraining (but not scruffing) the animals and manually attaching them to a fiber optic patch cord several times. Finally, if animals were included in DREADD manipulation cohorts, animals were habituated to intraperitoneal (IP) injections with saline every day of experimentation. After handling, animals were habituated to the empty arena for two days for 25 minutes/day. During habituation, animals were placed into the arena from their cage and allowed to explore the arena spontaneously. Finally, on the novelty session days, a single novel object (a single Mega Blok, Mega Bloks First Builders 80-piece Classic Building Bag) was briefly submerged in soiled bedding mixed from each mouse’s cage in the current round and placed in the corner of the arena (∼15 cm from either wall) (Akiti et al., 2022; Wooden et al., 2021). Animals were allowed to explore the arena as in habituation sessions for ∼25 min. Behavior was recorded with an overhead camera (Kinect v2, Microsoft for Motion Sequencing; FL3-U3-13E4M, PointGrey for all other experiments) at 15 or 30 Hz (15 Hz for Motion Sequencing Experiments; 30 Hz for all other experiments). Video data was captured for MoSeq using publicly available recording software; all other video data was recorded using Bonsai-rx (Lopes et al., 2015). Objects were cleaned with 70% ethanol at the end of each day and left out to dry overnight. The arena was cleaned with enzymatic cleaner (Nature’s Miracle) between animals every day and feces were removed. One object was used per animal and not reused across animals or cohorts. Animals were placed in the arena in the same order each day. For experiments with a feeding analysis, on the first novelty session day, after animals were removed from the arena, they were transferred to a recovery cage and given *ad libitum* food access. The amount the animal ate after three hours of food access was then recorded as the feeding amount for animals.

### Motion Sequencing

Motion Sequencing (MoSeq) was implemented as described previously (Wiltschko et al., 2015; Akiti et al., 2022). The analysis steps for MoSeq are briefly as follows. The raw imaging data is pre-processed with filtering and background subtraction, and the animal’s outline and position were extracted. Principal components (PC) are then calculated to represent the movements of each mouse to reduce the dimensionality of the behavioral data. At this step, 600 frames were trimmed from the start of each video to exclude timepoints when the animal is not in the box. This data in PC space is then modeled using a non-robust, single transition matrix, auto-regressive hidden Markov model (AR-HMM) to segment the continuous behavior into individual syllables. We used MoSeq to divide the data into 100 syllables used across days without the object present (two sessions) and with the object present (two sessions). All code for extracting and modeling the data are available using the pipeline found here: https://dattalab.github.io/moseq2-website/index.html. The syllables that explained a cumulative 90% of the total frames across all the sessions were used (Gschwind et al., 2023; Rudolph et al., 2020; Wiltschko et al., 2020), to a total of 58 syllables, for all analysis. Syllable ID numbers are arbitrary. Data from all behavioral sessions were used for syllable identification and initial modeling, but for post-hoc analysis, we split sessions based on whether an object was absent (days H1-H2) or present (days N1-N2).

### Syllable frequency analysis

Individual syllable frequencies were calculated across mice and across sessions for each type of session (empty arena, arena with object.) Syllable frequencies were normalized to type of session and syllables that are significantly enriched in one condition versus another (hungry versus sated animals) were identified using a Mann Whitney U test with Bonferroni corrections for multiple comparisons.

### Entropy analysis

Entropy was calculated using the Shannon entropy formula (H =−Σ_i_𝜋𝜋_𝑖𝑖_log (𝜋𝜋_𝑖𝑖_)). Syllable transition entropy was calculated by generating a bigram transition matrix for each animal and session. Entropy was then calculated on this entire transition distribution, to model the randomness of the transition distribution on all transitions (including self-transitions). Entropy was calculated per session and animal and then averaged within animal and type of session (e.g. empty arena vs. arena with object present.) For rolling entropy analysis, a single session per animal was divided into quintiles and a separate transition matrix was built for each quintile and entropy calculations were made for those quintiles.

### DeepLabCut analysis

To track the animal’s position relative to the object as well as quantify interactions with the object, we used DeepLabCut (Mathis et al., 2018) for pose estimation. We trained two separate networks for animals with fiber implants and for animals without fiber implants. We manually annotated the animal’s nose, head, and base of tail and trained a network on 20% of labeled frames. After using DeepLabCut to track these bodyparts in each video, we processed the files by first trimming those frames with <90% likelihood value, interpolating the animal’s position in between these frames, and smoothing the trajectory of the body part initially with a five-frame moving median filter (as recommended in DeepLabCut) and then post hoc with a 15-frame moving average filter. We defined the “interacting radius” as when one of the three body parts was within a 7 cm radius of the object. The beginning of a bout (start of approach) was defined as the earliest time of interactions to the time at which either the nose or the tail was within the interaction radius of the object. During each interacting timepoint, we determined whether the nose or tail was closer to the center of the object using the Euclidean distance. The end of a bout was defined as the time at which neither nose nor tail were within the interacting radius of the object (end of retreat). The start of retreat was set to be the timepoint when the nose was closest to the object during an interacting bout. The above-mentioned metrics of interacting radius as well as timepoints for exploration were based on prior studies (Akiti et al., 2022; Menegas et al., 2018).

### Stereotaxic surgical procedures

For all procedures, mice were anesthetized with inhaled isoflurane (1-3%) throughout the procedure and given *ad libitum* access to oral carprofen for a day before the operation. Under the stereotaxic frame (David Kopf Instruments), the skull was exposed and leveled and a small burr hole was drilled. AAVs that encoded DREADDs that required Cre for expression were injected initially in animals that did not express Cre recombinase to determine a concentration that showed minimal Cre-independent recombination and subsequent expression at the injection site. After burr hole creation, virus was injected at a rate of either 50 nL/min (TS) or 70 nL/min (SNc/SNpl, ARC) with a syringe pump (Harvard Apparatus, #883015). Pipettes were then allowed to rest at the injection site and above the injection site for at least 5 minutes each before they were slowly withdrawn (< 10 um/sec) from the injection site. The following volumes (in nL) and coordinates were used for each site (represented as mm from bregma in the order AP, ML, DV):

TOS: 300 nL at -1.4, 3.28, -2.45

VLS: 100 nL at 0.5, 3.4, -3.25 with a 15° angle

ARC: 300 nL at -1.6, ±0.26, -5.85

SNc: 250 nL at -3.0, ±1.4, -4.0

SNpl: 250 nL at -3.1, ±2.1, -3.6

For AAV injections alone, the skin was then sutured shut. For photometry recordings and optogenetic manipulations, the skull was then lightly scored with a razor blade to ensure adequate adhesion of implants. Animals were implanted with a fiber optic (MFC_200/230-0.37_3mm_MF1.25_FLT; Doric Lenses). For photometry recordings in particular, the implant was lowered to a depth of 150 µm above the virus injection site. The fiber was held in place with superglue (Loctite gel #454) and Metabond dental cement. For photometry experiments, recordings began at least three weeks post operation. For DREADD and optogenetic manipulation experiments, manipulations and recordings began at least four weeks post operation.

AAVs and the concentrations (in gc/mL) used in this study were as follows:

AAV9-CAG-dLight3.8: 1.0 x 10^13^ (UNC Neurotools)

AAV9-hSyn-dLight3.8: 2.1 x 10^12^ (UNC Neurotools)

AAV9-hSyn-dLight3.8mut: 2.31 x 10^13^ (Plasmid design/modification made in-house, mutation and cloning done by Azenta Genewiz, packaged at UNC Neurotools. Plasmid available on Addgene)

AAV9-hSyn-mCherry: 1.1 x 10^12^ (Addgene #114472)

AAV9-hSyn-DIO-hM3D(G)q-mCherry: 1.1 x 10^13^ (Addgene #44361)

AAV8-hSyn-DIO-hM4D(G)i-mCherry: 2.1 x 10^13^ (Addgene #44362)

AAV8-hSyn-DIO-mCherry: 2.1 x 10^12^ (Addgene #50459)

AAV8-hSyn-FLEX-TC66T-P2A-EGFP-P2A-N2CG: 1.2 x 10^12^ (Plasmid designed in-house, cloned by Epoch Life Science and Azenta Genewiz, packaged at UNC Neurotools. Plasmid available on Addgene.)

AAV8-hSyn-FLEX-TVA-EGFP: 1.0 x 10^13^ (Received under MTA from Salk Institute) AAVDJ/9-nEF-Con/Foff 2.0-ChRmine-oScarlet: 8.7 x 10^12^ (Plasmid from Addgene #137161 and packaged at Janelia Viral Core)

AAVDJ/9-hSyn-FLEX-10xmyc: 5.0 x 10^12^ (Plasmid produced by Twist BioScience as in Capelli et al., 2017 and packaged at Janelia Viral Core)

AAVDJ/9-hSyn-FLEX-H2B-GFP: 1 x 10^13^ (Plasmid produced by Twist BioScience as in Ruder et al., 2021 and packaged at Janelia Viral Core)

### Fiber photometry (intensity) recording

For intensity fiber photometry experiments, a fiber optic implant on each animal was connected with a 0.37 NA, 3 meter long low autofluorescence patchcord (MFP_200/220/900-0.37_3m_FCM-MF1.25, Doric Lenses) to a Doric minicube with integrated photodetector (iFMC5-G2_E1(460-490)_F1(500-540)_E2(555-570)_F2(580-680)_S, Doric Lenses). Excitation light was generated from blue (470-nm excitation light, M470F3, Thorlabs; LED driver LEDD1B, Thorlabs) and green (470-nm excitation light, M565F3, Thorlabs; LED driver LEDD1B, Thorlabs) LEDs modulated at 200 and 250 Hz, respectively. Excitation light was maintained at an average of 15-30 uW for blue and green light. Signals were amplified in DC mode in the integrated Doric minicube and collected by a data acquisition board (PCI-6115, National Instruments) at 4 kHz, controlled by ScanImage in MATLAB 2012a. The acquisition board also received a synchronizing pulse generated by the behavior system for alignment of behavioral and neural time series data.

### Fiber photometry (intensity) signal analysis

Intensity fiber photometry signals were analyzed as described previously (Chantranupong et al., 2022). Briefly, raw signals were first detrended using a rolling Z-score with a time window of 30 seconds (12000 samples). This initial detrending reduced artifacts of photobleaching of the fluorophore over the entire session.

Detrended, frequency-modulated signal was then demodulated using the Python spectrogram function implemented from the scipy package, with a window of 216 samples with an overlap of 108 samples, corresponding to a final sampling period of 27 ms. The power of the signal at the frequency band closest to the carrier frequency was taken as the demodulated signal. Finally, to quantify fluorescent transients as Z-scores, the signal was passed through an additional 30 second Z-score window. To synchronize this neural time series data with the behavioral data, a synchronizing pulse was sent from the Bonsai system recording the video to the data acquisition board logging neural time series data at the beginning and end of the session. Timepoints for each frame were interpolated within these synchronizing pulses and the closest timepoint in the demodulated (downsampled) photometry data was taken for each behavioral timepoint used for aligning and averaging photometry signals (approach start, retreat start, and retreat end). For comparisons of photometry signal between groups of animals, signals were averaged within each animal across interaction bouts, and then across animals within a group. To compare the retreat-associated TOS DA signal between groups, we analyzed the average Z-score from 1 second before to 1 second after retreat start. We used a broad window to capture the entire magnitude of the TOS DA transient when animals retreat from the novel object.

### Fluorescence lifetime photometry (FLiP) recordings

Fluorescence lifetime photometry was performed using a custom-built high-speed system that will be described in detail in an upcoming manuscript (Lodder *et al.,* In Preparation). Animals were connected with a patch cord (MFP_200/220/900- 0.37_2m_FCM-MF1.25, Doric Lenses) to the above-described system and allowed to habituate to the patch cord, with the laser on, in their home cage, for two to three minutes. The light intensity was matched across animals at 12 µW.

### FLiP signal analysis

For comparisons of photometry signal between groups of animals, signals were averaged within each animal across interaction bouts, and then across animals within a group. To compare the retreat-associated TOS DA signal between, we took the average lifetime signal from 1 second before to 1 second after the start of retreat (the same window used as in intensity fiber photometry signal analysis).

### DREADD manipulations of AgRP neurons during novel object exploration

For cohorts in which we employed DREADD manipulations of AgRP neurons, the novel object behavioral assay differed from the above section. For these animals, alongside the normal handling and habituation, on all days leading up to the novelty session day, they were given an intraperitoneal (IP) injection of 0.9% sterile saline to habituate the animals to the injection itself. After injection, on arena habituation days, the animals were transferred back to their home cage for 30 minutes and then placed into the arena. On the novelty session day, all animals (control and DREADD animals) were injected IP with 0.2 mg/kg of deschloroclozapine (DCZ) and then transferred to a recovery cage without food present. After 30 minutes, animals were transferred to the arena with the novel object present. After the novel object assay, animals were transferred to a feeding cage as described above, to measure feeding changes induced by AgRP manipulation.

### Optogenetic calibration of dopamine release in the tail of striatum

To calibrate the optogenetic stimulation of TOS-projecting DANs, animals were first exposed to ten 0.5 mA, 500 ms long, uncued footshocks in a white plexiglass box with a grid floor through which shocks were delivered (Med Associates #ENV-005A). Shocks were delivered through the grid floor spaced apart by 20-30 seconds (with the intershock interval randomly chosen from a uniform distribution.) We then, for each site, tested a number of optogenetic stimulation parameters by varying the frequency, pulse width, and power of optogenetic stimulation train, though each stimulation train was for a duration of 500 ms (to best match the kinetics of release evoked by a 500 ms footshock). A 500 ms pulse train with 5 ms pulse width of 5 mW delivered at 15 Hz was chosen as best matching the DA release profile in the TOS caused by a footshock.

Animals that were used for calibration of the optogenetic stimulation were not used in the novel object exploration behavioral task.

### Optogenetic manipulations of dopamine release during novel object exploration

For cohorts in which we employed optogenetic manipulations of DANs, animals were handled and habituated to the arena as in photometry recordings. On the novel object exposure day, Bonsai-rx was used for closed loop optogenetic stimulation. In Bonsai, a roughly circular ROI was drawn around the object ∼7 cm in radius (the same interacting radius as used for defining interacting time in DeepLabCut). The image was then thresholded such that the dark animal appeared white and the background (floor, walls, object etc.) all appeared black. To prevent the patch cord from being detected as the animal in the ROI, white tape was wrapped around the patchcord. The ROI was then calibrated such that if any part of the animal’s body fell within the defined ROI, a 1 Hz pulse was generated from an Arduino Uno. This 1 Hz pulse was sent to a Master 8 pulse generator which then generated a pulse pattern for optogenetic stimulation (500 ms pulse train consisting of a 15 Hz pattern of 5 ms pulses at 5 mW per pulse). This resulted in a 50% duty cycle of stimulation, to evoke individual DA pulses rather than a sustained increased baseline DA with continuous pulsed stimulation. This optogenetic stimulation parameter was fed into an acousto-optic modulator (AOM) placed after a 635 nm red laser (OptoEngine). An AOM was used instead of directly modulating the laser using a BNC port since the AOM allows for noiseless sub-millisecond modulation of laser power with minimal ramping. A shutter was not preferred since loud noises are known to induce DA release in the TOS (Menegas et al., 2018; Tsutsui-Kimura et al., 2022). After the novel object assay, animals were transferred to a feeding cage to assess feeding behavior as described above. After assessment of feeding behavior, animals were transferred to the FLiP system and the evoked change in lifetime in the TOS due to optogenetic stimulation was measured for each side separately. The peak lifetime evoked by stimulation was calculated for both sides of the brain for an animal and averaged. Post-hoc verification of the signal was preferred over real time validation of TOS DA release given difficulties in attaching more than two patch cords to the animal without hindering mobility.

### Rabies tracing

For rabies tracing experiments, slight modifications were made to the surgical procedure outlined above. Animals were first injected with a Cre-dependent AAV either with a bicistronic construct encoding both a weakened TVA (TC66T) and G protein (N2C(G)) or a Cre-dependent AAV with TVA-GFP alone in the SNc and SNpl. A burr hole was made in the same surgery over the TOS, and the surgical site was closed with sutures. After waiting 4 weeks to allow for sufficient expression of the helper proteins, the surgical site was reopened and EnvA-pseudotyped CVS-N2C-RbV-ΔG-H2b- tdTomato (4E9 pg/mL) was injected in the TOS. After waiting six days to allow for the rabies to infect axons and move retrograde transsynaptically, animals were perfused and histological sections were taken, as described below.

### Histology and Immunohistochemistry

Mice were anesthetized with higher concentration isoflurane inhalation and transcardially perfused with phosphate-buffered saline (PBS) and subsequently 4% paraformaldehyde (PFA) in PBS. Brains were then extracted and stored in 4% PFA/PBS overnight and then transferred to PBS for long-term storage. Brains were sliced using a Leica VT1000s vibratome into 70 µm thick free-floating sections. Slices were transferred to a six well plate and washed with PBS. Slices were then blocked at room temperature for two hours in 5% normal goat serum (Abcam), 0.1% TritonX-100 PBS. Slices were then placed into a solution with primary antibody (all at 1:1000) and allowed to incubate overnight at 4 °C. The following day, slices were washed with PBS with 0.1% TritonX-100 (PBST). Slices were then transferred to a solution with secondary antibody at 1:500 in blocking buffer. Slices were washed finally with PBST and placed in PBS before being mounted on a slide for imaging with ProLong Diamond Antifade Mountant with DAPI (ThermoFisher Scientific). Slices were imaged with an Olympus VS200 slide scanning microscope or for high-resolution images, a Yokagawa CSU-X1 spinning disk confocal microscope.

### Data Analysis/Statistics

All data was analyzed using Python v3.8. No statistical method was used to pre- determine sample size, though we attempted to match sample size across experiments. For DREADD and optogenetic manipulation experiments, experimenters were blinded to the animal condition. Experimenters were not blinded to the animal’s food restriction status given the need for placing special cage cards designating food restriction status. Non-parametric, two-sided statistical tests were used for all analysis and statistical significance was taken as p < 0.05. When necessary, Bonferroni correction for multiple comparisons was employed. In figures, n/s:p>.05, *:p<.05, **:p<.01, ***:p<.001. Data was not systematically tested for normalcy, hence the use of non-parametric statistics. All bar graphs are shown as the mean with error bars as the standard error about the mean (SEM). For all experiments, if an animal displaced the block from the floor of the arena, the data was not used (n=1). For refeeding experiments, animals that removed the food out of the food bowl, and thus it was difficult to measure amount eaten, were not included in feeding analysis but were included in the rest of the behavioral analysis (n=1).

## Acknowledgements

We thank Allison Girasole, Shun Li, Shijia Liu, Bastijn van den Boom, and Daniel Hochbaum of the Sabatini lab for helpful conversations throughout this process. We thank Adam Granger for designing the AAV8-hSyn-FLEX-TC66T-P2A-EGFP-P2A-

N2CG construct. We thank Seul Ah Kim for assistance in setting up the footshock apparatus. We thank Ofer Mazor and Pavel Gorelik of the Harvard Medical School Research Instrumentation Core for assistance in setting up the novel object exploration assay. Diagrams with pictures of a mouse were created using Biorender. This work is supported by grants to B.L.S. (NIH U19NS113201, R37NS046579, Howard Hughes Medical Institute), to T.K. (NIH F30DA061675), to I.G. (NIH F30MH132298), to N.U. (NIH U19NS113201, R01MH133950), and to M.W.U. (R01MH125162, R01MH133950, R01DA059751).

## Author Contributions

T.K., J.B.W., and B.L.S designed all experiments. T.K, E.B., M.R., J.L., and L.H. collected all data. B.L. built the FLiP system. P.C. validated the AAVDJ/9 construct. I.G. assisted in MoSeq setup. L.T. and R.D. designed and provided the dLight3.8 sensor.

M.W.U. and N.U. provided advice on dopamine signaling. All authors edited the manuscript.

---

**Figure S1:**
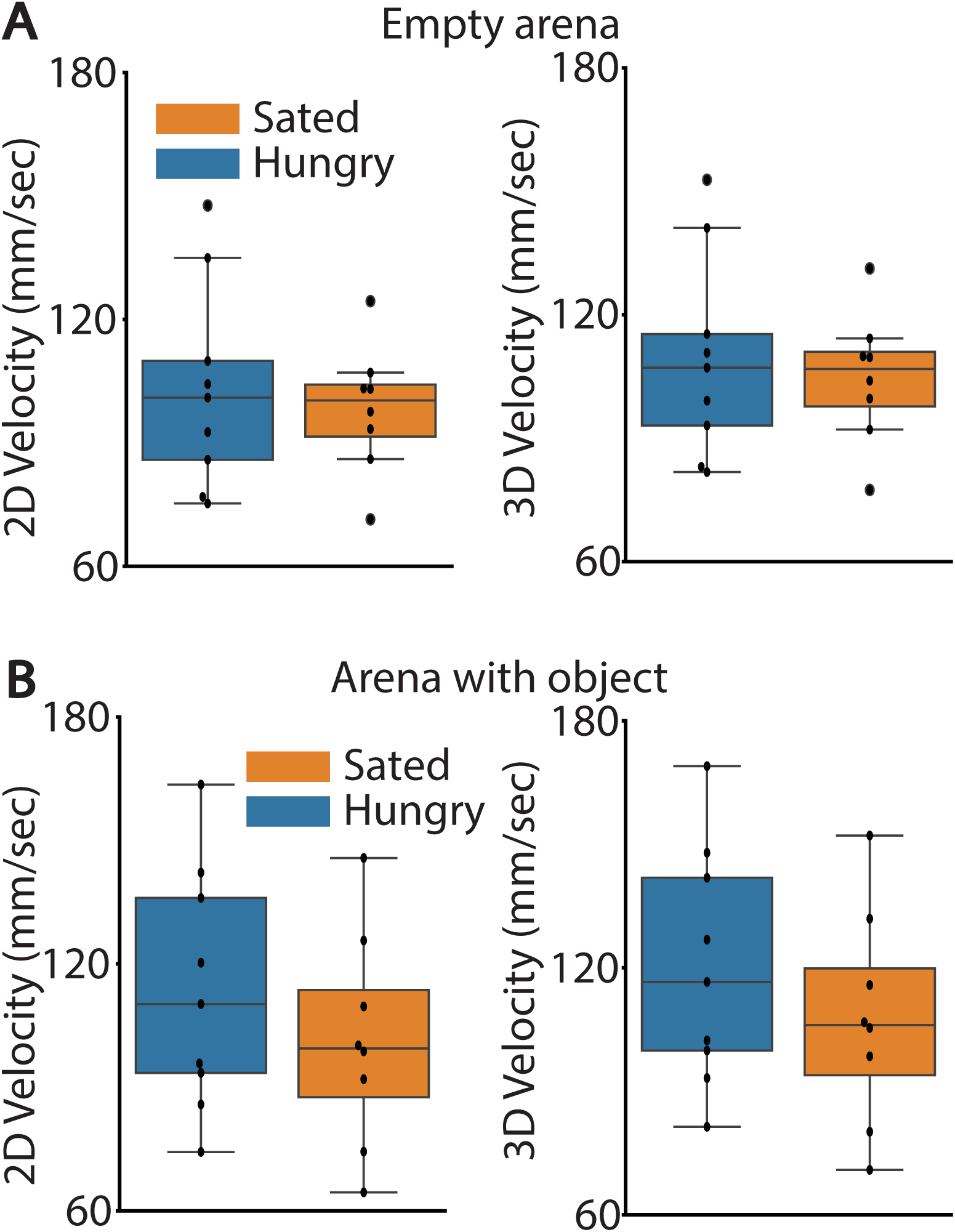
Hunger modulation of gross kinematic parameters and open field exploration. **A)** Comparison of two-dimensional (left; U=38, p=0.888) and three-dimensional (right; U=39, p=0.815) velocity of hungry and sated mice, averaged within mouse across only habituation sessions. **B)** As S1A-B, except for days with the novel object present (two-dimensional: U=43, p=0.541; three-dimensional: U=44, p=0.481)

**Figure S2:**
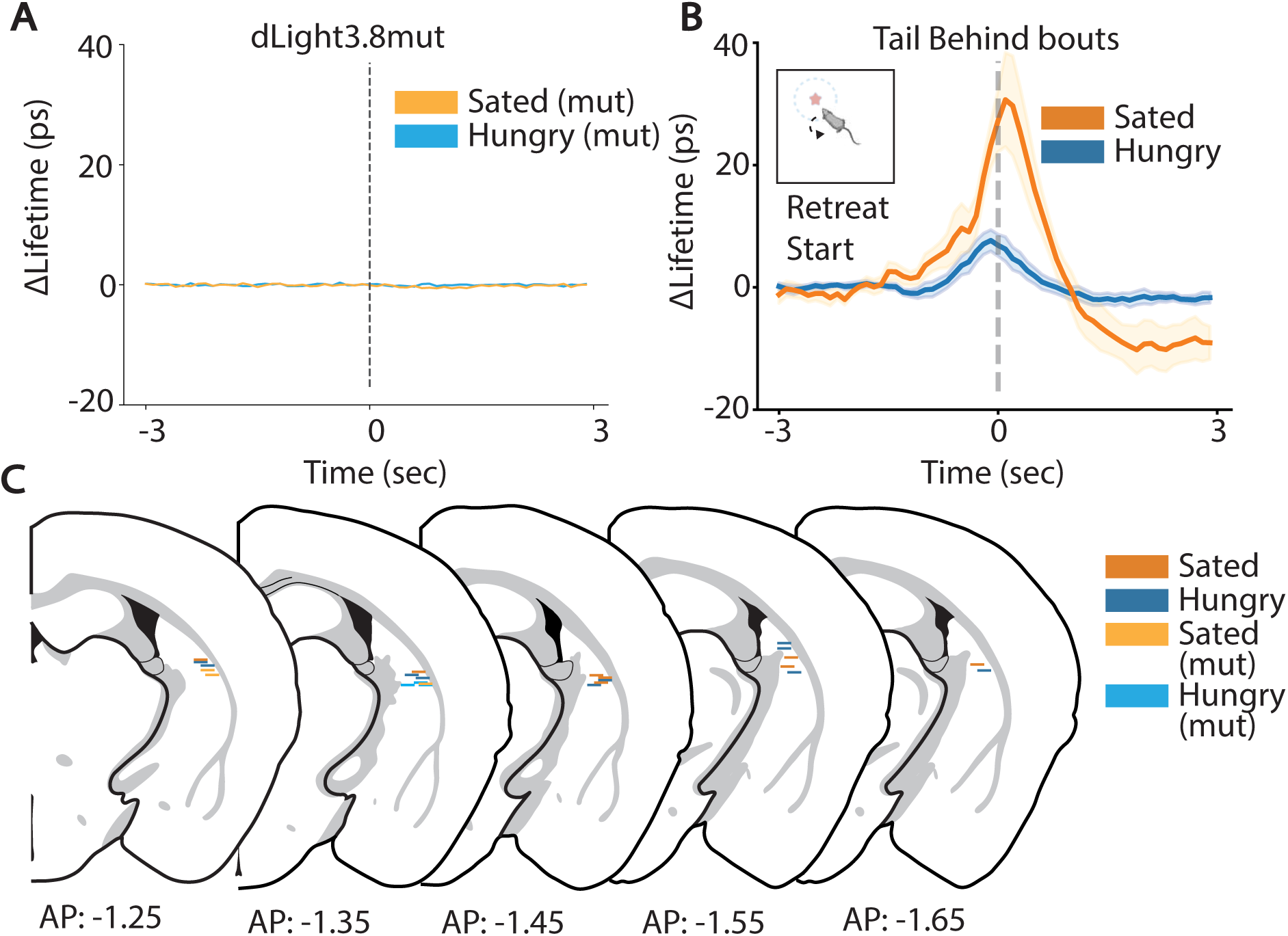
Schematic of fiber placements in TOS for Figure 3, and mutant sensor data. **A)** Comparison of changes in dLight3.8mut lifetime signal in hungry and sated animals retreating from a novel object. **B)** Comparison of dLight3.8 lifetime signal on trials in which both hungry and sated animals retreated from the object with tail behind the entire time. Bold line represents mean and shaded area is SEM calculated across animals. **C)** Schematic of fiber tip placements for dLight3.8 lifetime recordings in the TOS. Each bar represents a fiber optic tip.

**Figure S3:**
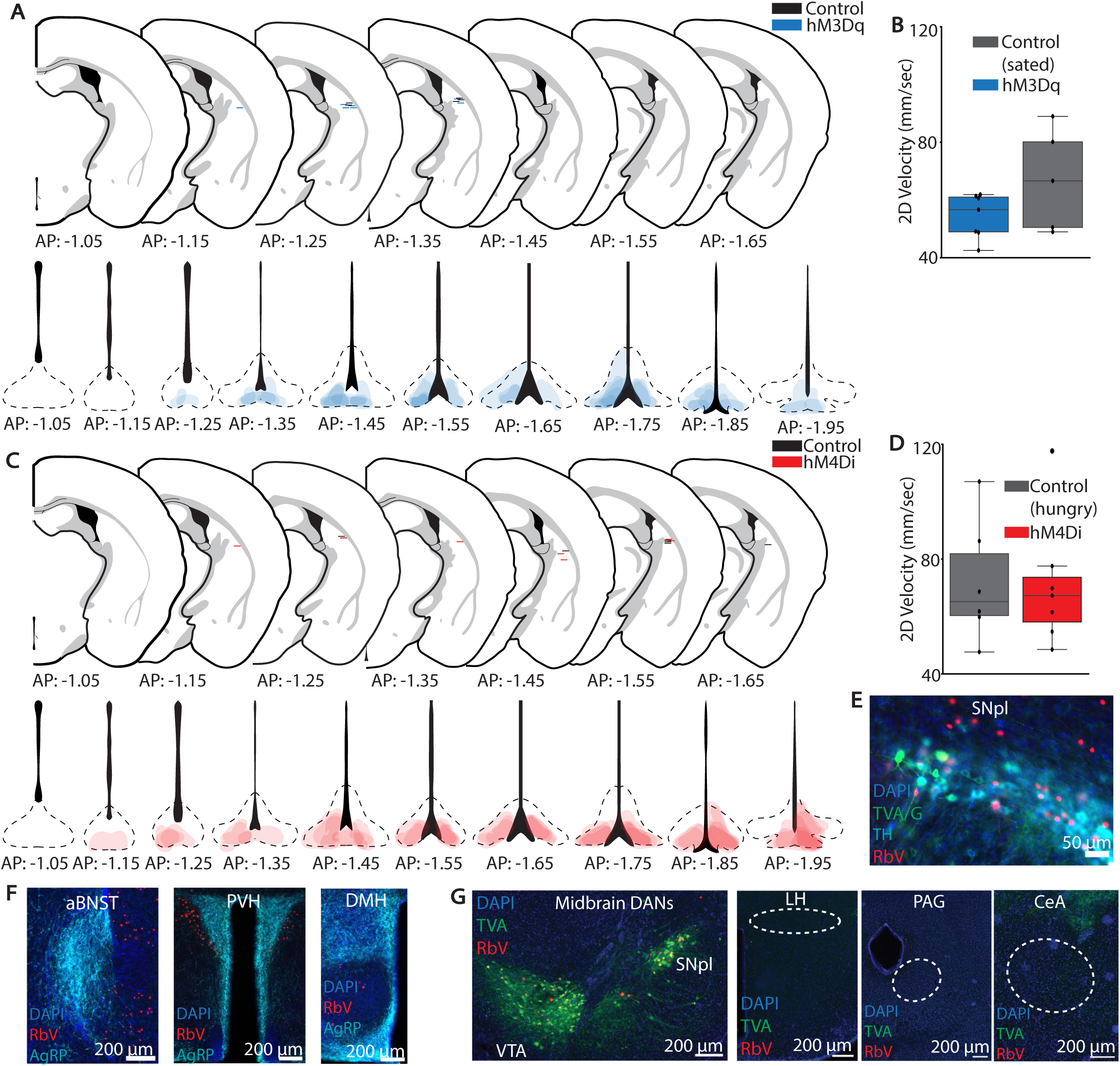
Schematic of DREADD AAV viral spread and photometry fiber tip locations in ARC and TOS, respectively and extended rabies tracing results, for Figure 4. **A)** TOS fiber placements and contour map of hM3Dq viral spread **B)** Comparison of two-dimensional velocity of hM3Dq versus control sated mice **C)** TOS fiber placements and contour map of hM4Di viral spread **D)** Comparison of two-dimensional velocity of hM4Di versus control hungry mice **E)** Lack of significant/only minor overlap between AgRP axons and RbV+ neurons in other major AgRP projection fields (anterior bed nucleus stria terminalis, aBNST; paraventricular hypothalamus, PVH; dorsomedial hypothalamus, DMH) **F)** RbV+ neurons (red) at midbrain DANs of DAT-*ires-*Cre animals injected with cre-dependent AAV encoding TVA alone (green) (left) and lack of RbV+ neurons at LH, CeA, and PAG in same animal. Dashed white lines represent areas of AgRP+ axons and RbV+ neurons in animals that have both the TVA and G protein present (see Figure 4H).

**Figure S4:**
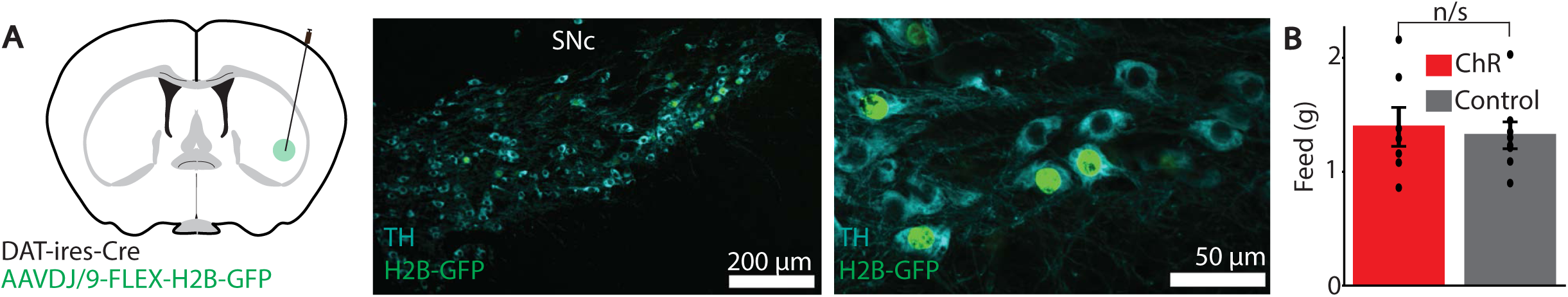
Validation of AAVDJ/9 infection of TH+ neurons and feeding analysis for optogenetic stimulation of TOS-projecting DANs, respectively for Figure 5. **A)** Injection schematic (injection of AAVDJ/9 virus in the ventrolateral striatum, VLS) and histological confirmation of AAVDJ/9 mediated retrograde infection of TH+ neurons in the midbrain. **B)** Comparison of feeding between animals transfected with opsin (ChRmine) and animals transfected with control non-opsin protein (10xmyc).

## Notes

### Competing Interest Statement

The authors have declared no competing interest.

## References

Akiti, K., Tsutsui-Kimura, I., Xie, Y., Mathis, A., Markowitz, J.E., Anyoha, R., Datta, S.R., Mathis, M.W., Uchida, N., Watabe-Uchida, M., 2022. Striatal dopamine explains novelty-induced behavioral dynamics and individual variability in threat prediction. Neuron 0. 10.1016/j.neuron.2022.08.022

Alhadeff, A.L., Goldstein, N., Park, O., Klima, M.L., Vargas, A., Betley, J.N., 2019.

Natural and Drug Rewards Engage Distinct Pathways that Converge on Coordinated Hypothalamic and Reward Circuits. Neuron 103, 891–908.e6. 10.1016/j.neuron.2019.05.050

Alhadeff, A.L., Su, Z., Hernandez, E., Klima, M.L., Phillips, S.Z., Holland, R.A., Guo, C., Hantman, A.W., Jonghe, B.C.D., Betley, J.N., 2018. A Neural Circuit for the Suppression of Pain by a Competing Need State. Cell 173, 140–152.e15. 10.1016/j.cell.2018.02.057

Aponte, Y., Atasoy, D., Sternson, S.M., 2011. AGRP neurons are sufficient to orchestrate feeding behavior rapidly and without training. Nat. Neurosci. 14, 351–355. 10.1038/nn.2739

Bäckman, C.M., Malik, N., Zhang, Y., Shan, L., Grinberg, A., Hoffer, B.J., Westphal, H., Tomac, A.C., 2006. Characterization of a mouse strain expressing Cre recombinase from the 3′ untranslated region of the dopamine transporter locus. genesis 44, 383–390. 10.1002/dvg.20228

Betley, J.N., Xu, S., Cao, Z.F.H., Gong, R., Magnus, C.J., Yu, Y., Sternson, S.M., 2015. Neurons for hunger and thirst transmit a negative-valence teaching signal. Nature 521, 180–185. 10.1038/nature14416

Bromberg-Martin, E.S., Hikosaka, O., 2011. Lateral habenula neurons signal errors in the prediction of reward information. Nat. Neurosci. 14, 1209–1216. 10.1038/nn.2902

Burnett, C.J., Li, C., Webber, E., Tsaousidou, E., Xue, S.Y., Brüning, J.C., Krashes, M.J., 2016. Hunger-Driven Motivational State Competition. Neuron 92, 187–201. 10.1016/j.neuron.2016.08.032

Capelli, P., Pivetta, C., Soledad Esposito, M., Arber, S., 2017. Locomotor speed control circuits in the caudal brainstem. Nature 551, 373–377. 10.1038/nature24064

Chantranupong, L., Beron, C.C., Zimmer, J.A., Wen, M.J., Wang, W., Sabatini, B.L., 2022. Local and long-distance inputs dynamically regulate striatal acetylcholine during decision making. 10.1101/2022.09.09.507130

Coddington, L.T., Lindo, S.E., Dudman, J.T., 2023. Mesolimbic dopamine adapts the rate of learning from action. Nature 614, 294–302. 10.1038/s41586-022-05614-z

Cone, J.J., McCutcheon, J.E., Roitman, M.F., 2014. Ghrelin Acts as an Interface between Physiological State and Phasic Dopamine Signaling. J. Neurosci. 34, 4905. 10.1523/JNEUROSCI.4404-13.2014

Corey, D.T., 1978. The determinants of exploration and neophobia. Neurosci. Biobehav Rev. 2, 235–253. 10.1016/0149-7634(78)90033-7

Dietrich, M.O., Zimmer, M.R., Bober, J., Horvath, T.L., 2015. Hypothalamic Agrp Neurons Drive Stereotypic Behaviors beyond Feeding. Cell 160, 1222–1232. 10.1016/j.cell.2015.02.024

Dodt, S., Widdershooven, N.V., Dreisow, M.-L., Weiher, L., Steuernagel, L., Wunderlich, F.T., Brüning, J.C., Fenselau, H., 2024. NPY-mediated synaptic plasticity in the extended amygdala prioritizes feeding during starvation. Nat. Commun. 15, 5439. 10.1038/s41467-024-49766-0

Douglass, A.M., Resch, J.M., Madara, J.C., Kucukdereli, H., Yizhar, O., Grama, A., Yamagata, M., Yang, Z., Lowell, B.B., 2023. Neural basis for fasting activation of the hypothalamic–pituitary–adrenal axis. Nature 620, 154–162. 10.1038/s41586-023-06358-0

Düring, D.N., Dittrich, F., Rocha, M.D., Tachibana, R.O., Mori, C., Okanoya, K., Boehringer, R., Ehret, B., Grewe, B.F., Gerber, S., Ma, S., Rauch, M., Paterna, J.-C., Kasper, R., Gahr, M., Hahnloser, R.H.R., 2020. Fast Retrograde Access to Projection Neuron Circuits Underlying Vocal Learning in Songbirds. Cell Rep. 33, 108364. 10.1016/j.celrep.2020.108364

Green, I., Amo, R., Watabe-Uchida, M., 2024. Shifting attention to orient or avoid: a unifying account of the tail of the striatum and its dopaminergic inputs. Curr. Opin. Behav. Sci. 59, 101441. 10.1016/j.cobeha.2024.101441

Greenstreet, F., Vergara, H.M., Pati, S., Schwarz, L., Wisdom, M., Marbach, F., Johansson, Y., Rollik, L., Moskovitz, T., Clopath, C., Stephenson-Jones, M., 2022. Action prediction error: a value-free dopaminergic teaching signal that drives stable learning. 10.1101/2022.09.12.507572

Grove, J.C.R., Gray, L.A., La Santa Medina, N., Sivakumar, N., Ahn, J.S., Corpuz, T.V., Berke, J.D., Kreitzer, A.C., Knight, Z.A., 2022. Dopamine subsystems that track internal states. Nature 608, 374–380. 10.1038/s41586-022-04954-0

Gschwind, T., Zeine, A., Raikov, I., Markowitz, J.E., Gillis, W.F., Felong, S., Isom, L.L., Datta, S.R., Soltesz, I., 2023. Hidden behavioral fingerprints in epilepsy. Neuron 111, 1440–1452.e5. 10.1016/j.neuron.2023.02.003

Gunaydin, L.A., Grosenick, L., Finkelstein, J.C., Kauvar, I.V., Fenno, L.E., Adhikari, A., Lammel, S., Mirzabekov, J.J., Airan, R.D., Zalocusky, K.A., Tye, K.M., Anikeeva, P., Malenka, R.C., Deisseroth, K., 2014. Natural Neural Projection Dynamics Underlying Social Behavior. Cell 157, 1535–1551. 10.1016/j.cell.2014.05.017

Hochbaum, D.R., Hulshof, L., Urke, A., Wang, W., Dubinsky, A.C., Farnsworth, H.C., Hakim, R., Lin, S., Kleinberg, G., Robertson, K., Park, C., Solberg, A., Yang, Y., Baynard, C., Nadaf, N.M., Beron, C.C., Girasole, A.E., Chantranupong, L., Cortopassi, M.D., Prouty, S., Geistlinger, L., Banks, A.S., Scanlan, T.S., Datta, S.R., Greenberg, M.E., Boulting, G.L., Macosko, E.Z., Sabatini, B.L., 2024. Thyroid hormone remodels cortex to coordinate body-wide metabolism and exploration. Cell 187, 5679-5697.e23. 10.1016/j.cell.2024.07.041

Krashes, M.J., Koda, S., Ye, C., Rogan, S.C., Adams, A.C., Cusher, D.S., Maratos-Flier, E., Roth, B.L., Lowell, B.B., 2011. Rapid, reversible activation of AgRP neurons drives feeding behavior in mice. J. Clin. Invest. 121, 1424–1428. 10.1172/JCI46229

Krausz, T.A., Comrie, A.E., Kahn, A.E., Frank, L.M., Daw, N.D., Berke, J.D., 2023. Dual credit assignment processes underlie dopamine signals in a complex spatial environment. Neuron 111, 3465–3478.e7. 10.1016/j.neuron.2023.07.017

Lee, S.J., Lodder, B., Chen, Y., Patriarchi, T., Tian, L., Sabatini, B.L., 2021. Cell-type specific asynchronous modulation of PKA by dopamine in learning. Nature 590, 451–456. 10.1038/s41586-020-03050-5

Lin, S., Gillis, W.F., Weinreb, C., Zeine, A., Jones, S.C., Robinson, E.M., Markowitz, J., Datta, S.R., 2024. Characterizing the structure of mouse behavior using Motion Sequencing. Nat. Protoc. 1–50. 10.1038/s41596-024-01015-w

Liu, Q., Yang, X., Luo, M., Su, J., Zhong, J., Li, X., Chan, R.H.M., Wang, L., 2023. An iterative neural processing sequence orchestrates feeding. Neuron 0. 10.1016/j.neuron.2023.02.025

Livneh, Y., Ramesh, R.N., Burgess, C.R., Levandowski, K.M., Madara, J.C., Fenselau, H., Goldey, G.J., Diaz, V.E., Jikomes, N., Resch, J.M., Lowell, B.B., Andermann, M.L., 2017. Homeostatic circuits selectively gate food cue responses in insular cortex. Nature 546, 611–616. 10.1038/nature22375

Long, C., Lee, K., Yang, L., Dafalias, T., Wu, A.K., Masmanidis, S.C., 2024. Constraints on the subsecond modulation of striatal dynamics by physiological dopamine signaling. Nat. Neurosci. 27, 1977–1986. 10.1038/s41593-024-01699-z

Lopes, G., Bonacchi, N., Frazão, J., Neto, J.P., Atallah, B.V., Soares, S., Moreira, L., Matias, S., Itskov, P.M., Correia, P.A., Medina, R.E., Calcaterra, L., Dreosti, E., Paton, J.J., Kampff, A.R., 2015. Bonsai: an event-based framework for processing and controlling data streams. Front. Neuroinformatics 9.

Lutas, A., Kucukdereli, H., Alturkistani, O., Carty, C., Sugden, A.U., Fernando, K., Diaz, V., Flores-Maldonado, V., Andermann, M.L., 2019. State-specific gating of salient cues by midbrain dopaminergic input to basal amygdala. Nat. Neurosci. 22, 1820–1833. 10.1038/s41593-019-0506-0

Mandelblat-Cerf, Y., Ramesh, R.N., Burgess, C.R., Patella, P., Yang, Z., Lowell, B.B., Andermann, M.L., 2015. Arcuate hypothalamic AgRP and putative POMC neurons show opposite changes in spiking across multiple timescales. eLife 4, e07122. 10.7554/eLife.07122

Markowitz, J.E., Gillis, W.F., Beron, C.C., Neufeld, S.Q., Robertson, K., Bhagat, N.D., Peterson, R.E., Peterson, E., Hyun, M., Linderman, S.W., Sabatini, B.L., Datta, S.R., 2018. The Striatum Organizes 3D Behavior via Moment-to-Moment Action Selection. Cell 174, 44–58.e17. 10.1016/j.cell.2018.04.019

Markowitz, J.E., Gillis, W.F., Jay, M., Wood, J., Harris, R.W., Cieszkowski, R., Scott, R., Brann, D., Koveal, D., Kula, T., Weinreb, C., Osman, M.A.M., Pinto, S.R., Uchida, N., Linderman, S.W., Sabatini, B.L., Datta, S.R., 2023. Spontaneous behaviour is structured by reinforcement without explicit reward. Nature 614, 108–117. 10.1038/s41586-022-05611-2

Marshel, J.H., Kim, Y.S., Machado, T.A., Quirin, S., Benson, B., Kadmon, J., Raja, C., Chibukhchyan, A., Ramakrishnan, C., Inoue, M., Shane, J.C., McKnight, D.J., Yoshizawa, S., Kato, H.E., Ganguli, S., Deisseroth, K., 2019. Cortical layer– specific critical dynamics triggering perception. Science 365, eaaw5202. 10.1126/science.aaw5202

Mathis, A., Mamidanna, P., Cury, K.M., Abe, T., Murthy, V.N., Mathis, M.W., Bethge, M., 2018. DeepLabCut: markerless pose estimation of user-defined body parts with deep learning. Nat. Neurosci. 21, 1281–1289. 10.1038/s41593-018-0209-y

Menegas, W., Akiti, K., Amo, R., Uchida, N., Watabe-Uchida, M., 2018. Dopamine neurons projecting to the posterior striatum reinforce avoidance of threatening stimuli. Nat. Neurosci. 21, 1421–1430. 10.1038/s41593-018-0222-1

Menegas, W., Babayan, B.M., Uchida, N., Watabe-Uchida, M., 2017. Opposite initialization to novel cues in dopamine signaling in ventral and posterior striatum in mice. eLife 6, e21886. 10.7554/eLife.21886

Mikhael, J.G., Gershman, S.J., 2022. Impulsivity and risk-seeking as Bayesian inference under dopaminergic control. Neuropsychopharmacol. Off. Publ. Am. Coll. Neuropsychopharmacol. 47, 465–476. 10.1038/s41386-021-01125-z

Millard, S.J., Hoang, I.B., Sherwood, S., Taira, M., Reyes, V., Greer, Z., O’Connor, S.L., Wassum, K.M., James, M.H., Barker, D.J., Sharpe, M.J., 2024. Cognitive representations of intracranial self-stimulation of midbrain dopamine neurons depend on stimulation frequency. Nat. Neurosci. 27, 1253–1259. 10.1038/s41593-024-01643-1

Miyamichi, K., Shlomai-Fuchs, Y., Shu, M., Weissbourd, B.C., Luo, L., Mizrahi, A., 2013. Dissecting Local Circuits: Parvalbumin Interneurons Underlie Broad Feedback Control of Olfactory Bulb Output. Neuron 80, 1232–1245. 10.1016/j.neuron.2013.08.027

Müller, M., Wehner, R., 1994. The hidden spiral: systematic search and path integration in desert ants, Cataglyphis fortis. J. Comp. Physiol. A 175, 525–530. 10.1007/BF00199474

Padilla, S.L., Qiu, J., Soden, M.E., Sanz, E., Nestor, C.C., Barker, F.D., Quintana, A., Zweifel, L.S., Rønnekleiv, O.K., Kelly, M.J., Palmiter, R.D., 2016. Agouti-related peptide neural circuits mediate adaptive behaviors in the starved state. Nat. Neurosci. 19, 734–741. 10.1038/nn.4274

Reardon, T.R., Murray, A.J., Turi, G.F., Wirblich, C., Croce, K.R., Schnell, M.J., Jessell, T.M., Losonczy, A., 2016. Rabies Virus CVS-N2cΔG Strain Enhances Retrograde Synaptic Transfer and Neuronal Viability. Neuron 89, 711–724. 10.1016/j.neuron.2016.01.004

Ruder, L., Schina, R., Kanodia, H., Valencia-Garcia, S., Pivetta, C., Arber, S., 2021. A functional map for diverse forelimb actions within brainstem circuitry. Nature 590, 445–450. 10.1038/s41586-020-03080-z

Rudolph, S., Guo, C., Pashkovski, S.L., Osorno, T., Gillis, W.F., Krauss, J.M., Nyitrai, H., Flaquer, I., El-Rifai, M., Datta, S.R., Regehr, W.G., 2020. Cerebellum-Specific Deletion of the GABAA Receptor δ Subunit Leads to Sex-Specific Disruption of Behavior. Cell Rep. 33, 108338. 10.1016/j.celrep.2020.108338

Schmack, K., Bosc, M., Ott, T., Sturgill, J.F., Kepecs, A., 2021. Striatal dopamine mediates hallucination-like perception in mice. Science 372, eabf4740. 10.1126/science.abf4740

Simpson, E.H., Akam, T., Patriarchi, T., Blanco-Pozo, M., Burgeno, L.M., Mohebi, A., Cragg, S.J., Walton, M.E., 2024. Lights, fiber, action! A primer on in vivo fiber photometry. Neuron 112, 718–739. 10.1016/j.neuron.2023.11.016

Smith, N.K., Grueter, B.A., 2022. Hunger-driven adaptive prioritization of behavior. FEBS J. 289, 922–936. 10.1111/febs.15791

Tong, Q., Ye, C.-P., Jones, J.E., Elmquist, J.K., Lowell, B.B., 2008. Synaptic release of GABA by AgRP neurons is required for normal regulation of energy balance. Nat. Neurosci. 11, 998–1000. 10.1038/nn.2167

Tsutsui-Kimura, I., Uchida, N., Watabe-Uchida, M., 2022. Dynamical management of potential threats regulated by dopamine and direct- and indirect-pathway neurons in the tail of the striatum. 10.1101/2022.02.05.479267

Wiltschko, A.B., Johnson, M.J., Iurilli, G., Peterson, R.E., Katon, J.M., Pashkovski, S.L., Abraira, V.E., Adams, R.P., Datta, S.R., 2015. Mapping Sub-Second Structure in Mouse Behavior. Neuron 88, 1121–1135. 10.1016/j.neuron.2015.11.031

Wiltschko, A.B., Tsukahara, T., Zeine, A., Anyoha, R., Gillis, W.F., Markowitz, J.E., Peterson, R.E., Katon, J., Johnson, M.J., Datta, S.R., 2020. Revealing the structure of pharmacobehavioral space through motion sequencing. Nat. Neurosci. 23, 1433–1443. 10.1038/s41593-020-00706-3

Wooden, J.I., Spinetta, M.J., Nguyen, T., O’Leary, C.I., Leasure, J.L., 2021. A Sensitive Homecage-Based Novel Object Recognition Task for Rodents. Front. Behav. Neurosci. 15, 680042. 10.3389/fnbeh.2021.680042

Zafiri, D., Salinas-Hernández, X.I., Biasi, E.S.D., Rebelo, L., Duvarci, S., 2023. Dopamine Prediction Error Signaling in a Unique Nigrostriatal Circuit is Critical for Associative Fear Learning. 10.1101/2023.12.08.570564

